# Dual-inactivation of Regnase-1 and SOCS1 rewires exhausted CD8^+^ T cell fate to enhance anti-tumor functionality

**DOI:** 10.64898/2026.01.21.700812

**Authors:** Isabelle Le Mercier, David Monteiro, Katarina Halpin-Veszeleiova, Karrie Wong, Anne Dodson, Gustavo J. Martinez, Dominick Matos, Bashar Hamza, Ashish Yeri, Seamus McKenney, Sharon Lin, Nicole Protheroe, Alex Smashnov, Paul Dunbar, Mitali Ghose, Conor Calnan, Hugh Gannon, Somya Jain, Frank Thompson, Sophie Capobianco, Glenn J. Hanna, Eric Fagerberg, Sol Shenker, Sean Keegan, Gregory Kryukov, Jason Merkin, Ribhu Nayar, Caroline Bullock, Chris Wrocklage, Louise Cadzow, Frank Stegmeier, Marie-Andrée Forget, Chantale Bernatchez, John Gagnon, Fiona McHugh, Dipen Sangurdekar, Michael Schlabach, Micah J. Benson

## Abstract

The solid tumor microenvironment inhibits the functionality of tumor infiltrating T cells recognizing cognate tumor antigen, driving their differentiation towards terminal exhaustion. Interventions are sought to enhance the anti-tumor functionality of tumor-reactive T cells for clinical benefit. The functional genome regulating CD8^+^ T cell function against solid tumors was mapped by performing genome-wide, focused, and combination *in vivo* CRISPR/Cas9 screens using OT1 and PMEL TCR transgenic T cells in B16-OVA, MC38-gp100 and EG7-OVA syngeneic tumor models. The ability of the top single hits and combinations, which include Regnase-1 and SOCS1, to enhance CD8^+^ T cell anti-tumor function was evaluated in the OT1/B16-OVA model with large and established tumors, the disseminated PMEL/B16F10 tumor model, and in a novel murine TIL syngeneic model. The impact of Regnase-1 and SOCS1 single and dual-inactivation on the differentiation of exhausted CD8^+^ T cell subsets and on long-term persistent memory following tumor clearance was evaluated in OT1 CD8^+^ T cells in the B16-OVA model. The impact of single and dual-inactivation of Regnase-1 and SOCS1 on the anti-tumor function of experimental human T cell therapeutics was characterized in CRISPR/Cas9-engineered human TIL derived in vitro and in mesothelin-targeting CAR-Ts in vivo. NF-κB and cytokine signaling were identified as the top pathways regulating CD8^+^ T cell anti-tumor function, with Regnase-1 and Suppressor of Cytokine Signaling 1 (SOCS1) the top single and combination edits regulating the accumulation of tumor-specific TCR transgenic CD8^+^ T cells in syngeneic tumor models. Dual-inactivation of Regnase-1 and SOCS1 cooperated through non-redundant mechanisms to strongly expand intermediate (Tex^int^) and effector (Tex^eff^) exhausted CD8^+^ T cells within lymphoid tissues and tumor, with CD8^+^ T cells rewired to display an enhanced effector state and suppressed expression of TOX. Dual-edited persistent T effector memory cells (T_em_) were formed following tumor clearance. Lastly, Regnase-1 and SOCS1 inactivation enhanced human Tumor Infiltrating Lymphocyte (TIL) and chimeric antigen receptor T cells (CAR-T) therapy functionality. Collectively, this study systematically mapped pathways regulating CD8^+^ T cell anti-tumor functionality, with Regnase-1 and SOCS1 dual-inactivation found to maximize anti-tumor function through non-redundant mechanisms.

## BACKGROUND

T lymphocytes can directly recognize and kill tumors to drive meaningful clinical responses in patients with solid tumors[1,2]. Existing therapeutic interventions co-opt this mechanism and include the adoptive transfer of large numbers of tumor-targeting T cells into patients[3,4]. Unfortunately, most solid tumor patients fail to respond to T cell-driven immunotherapies due in part to the formidable barrier posed by the immunosuppressive tumor microenvironment[5]. In pre-clinical mouse models as well as patients, tumor-specific CD8^+^ T cells can be found in a T cell exhaustion (Tex) differentiation continuum[6–8] which includes stem-like Tex progenitors (Tex^prog^) possessing self-renewal capacity driven by transcription factor TCF1[7,9–12], proliferating intermediate (Tex^int^) cells with circulatory and effector capacity[13], effector (Tex^eff^) cells optimized for cytolytic killing of tumor[14–16] and terminally differentiated functionally exhausted cells (Tex^term^) driven by the transcription factor TOX[13,17,18]. Therapeutic interventions seeking to harness T cell anti-tumor functionality should ideally impact across Tex subsets.

Mapping intrinsic pathways restraining CD8^+^ T cell anti-tumor functionality is essential to rationally design new therapeutics, with CRISPR/Cas9 functional genomic screens serving as a powerful tool to define the therapeutic nodes regulating lymphocyte function and differentiation[19–23]. In vivo CD8^+^ T cell screens conducted to date have successfully identified novel genes yet have been limited by either library size or depth of sgRNA recovery, with a comprehensive map of the functional genome regulating CD8^+^ T cell anti-tumor function yet to be obtained[24–30]. Systematic efforts to identify the top dual-edit combinations maximizing in vivo CD8^+^ T cell anti-tumor function have additionally not been undertaken. We expanded on our previous focused in vivo CRISPR screen[31] efforts to perform genome-wide, focused and combination screens to map the functional genome of CD8^+^ T cell anti-tumor functionality. Negative regulators of NF-κB and cytokine signaling emerged as the top pathways, with Regnase-1 and SOCS1 the top single and dual-edits. Regnase-1 is a dynamically regulated RNA-binding protein with ribonuclease activity targeting mRNA substrates including *c-Rel, Icos* and *Batf* in T cells whose inactivation has been shown to enhance anti-tumor effector cell activity[28,32,33]. SOCS1 is a negative regulator of cytokine signaling[34] whose inactivation in CD8^+^ T cells we and others have shown also enhances anti-tumor activity[19,31] by expanding central memory (T_cm_) cells as well as tumor-infiltrating Tex^int^ and Tex^eff^ cells[31]. We demonstrate that dual-inactivation of Regnase-1 and SOCS1 maximizes CD8^+^ T cell anti-tumor functionality by perturbing the differentiation trajectory and functional status of Tex and memory subsets through complementary mechanisms and extend these insights to develop novel CRISPR/Cas9 engineered TIL (eTIL^®^) and CAR-T (eCAR^™^-T) therapeutics for the treatment of refractory solid tumors.

## RESULTS

### In vivo CRISPR/Cas9 screens map the CD8^+^ T cell functional genome to identify the NF-κB and cytokine signaling pathways as critical regulators of anti-tumor function

To map the genome for regulators of CD8^+^ T cell function against solid tumors, 21,422 genes (library design detailed in Supplementary Methods) were subdivided into 24 sgRNA library ‘bookshelves’ each targeting ∼900 genes, with 8 sgRNAs per gene. This bookshelf approach was used to ensure sufficient sgRNA coverage upon recovery, addressing the limitation of coverage when intact genome-wide libraries are used in vivo settings[29]. Unique Molecular Identifiers (UMIs) were additionally included to track sgRNAs, with the use of bookshelves together with UMIs intended to detect enriched and depleted sgRNAs at high sensitivity[27]. We crossed OVA peptide-recognizing OT1 TCR-Tg[35] mice with Cas9 transgenic mice[36], with Cas9-expressing OT1 TCR-Tg CD8^+^ T cells transduced with sgRNA bookshelves and transferred into B16-OVA tumor-bearing mice (Figure 1A). Comparison of the distribution of UMIs between input and tumor-infiltrating OT1s yielded genes positively or negatively influencing OT1 accumulation within tumor (Figure 1A, Supplementary Table S1). Depletion of essential genes from DepMap (Figure S1A) and cumulative recovery of 85% of total sgRNAs from tumors (Figure S1B) were measures of screen robustness, validating the bookshelf approach and verifying coverage of the genome. We used both lentiviral and retroviral vectors during screening, with concordant screen results obtained (Figure S1C). Informed by our previous results [31], each bookshelf contained positive and negative enrichment, non-cutting, and olfactory control sgRNAs [31].

**Figure 1:**
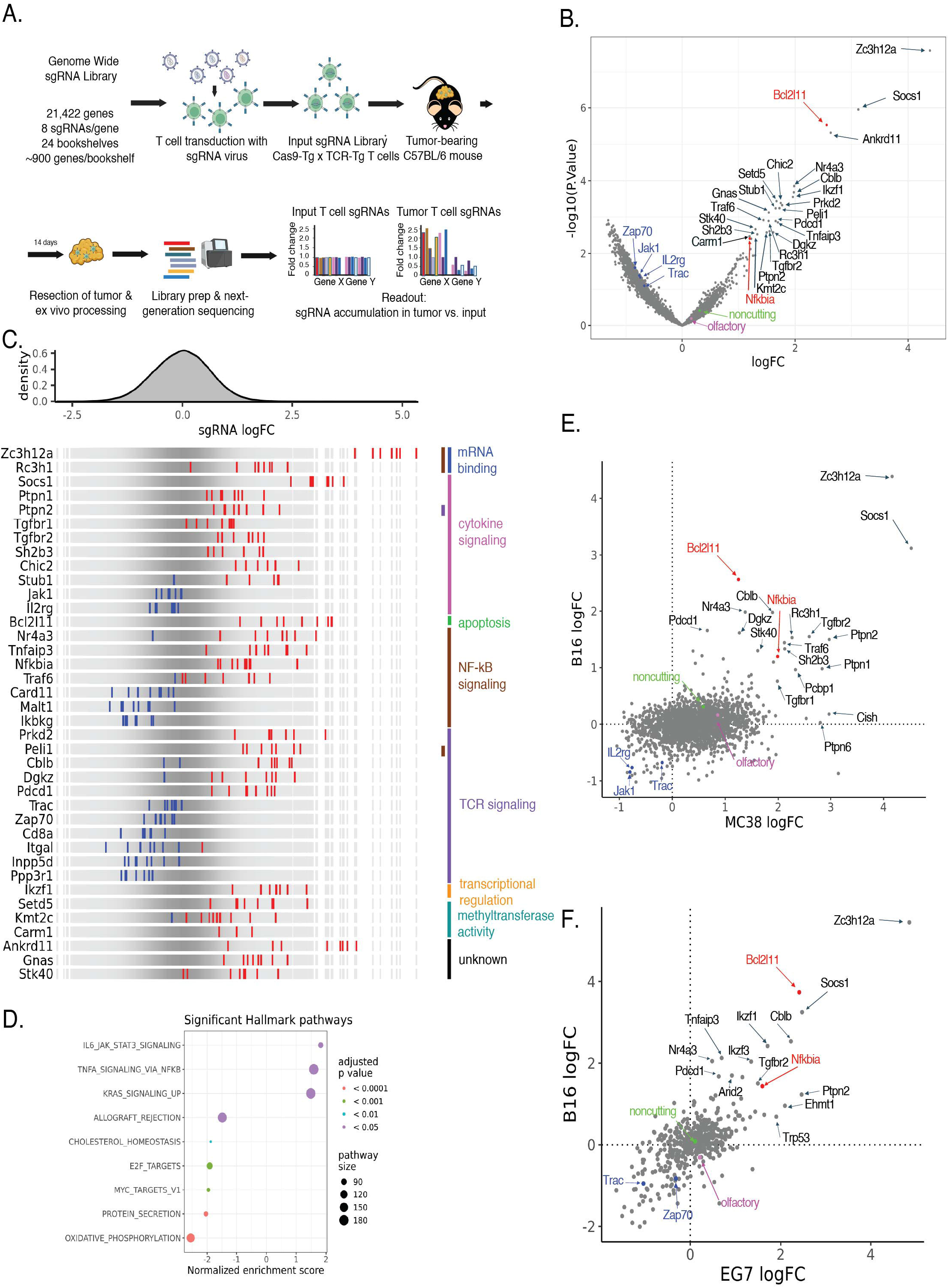
In vivo CRISPR/Cas9 screens map the CD8^+^ T cell functional genome to identify the NF-κB and cytokine signaling pathways as critical regulators of anti-tumor function. **(A)** Experimental schema depicting a genome-wide in vivo CRISPR screen using Cas9-Tg OT1 TCR-Tg CD8^+^ T cells. sgRNA^+^ OT1s were transferred into B16-OVA tumor-bearing mice, with tumors harvested 14 days following transfer, and the sgRNA distribution of OT1s from tumors and input analyzed. **(B)** Results of genome-wide screen identifying top hits whose inactivation enhances OT1 CD8^+^ T cell accumulation in tumors. Positive, negative and inert/noncutting controls are represented in red, blue, purple and green, respectively. **(C)** Enrichment (red) or depletion (blue) patterns of individual sgRNAs targeting gene hits and controls by tumor infiltrating OT1s. Genes are clustered by established protein function, as indicated. **(D)** Hallmark gene sets significantly enriched for hits shown in Figure 1B. **(E)** Results of an 8,006 gene screen identifying top hits enhancing PMEL CD8^+^ T cell accumulation into MC38-gp100 tumor bearing mice, with tumors harvested 14 days after transfer. Results are depicted in comparison to a OT1 / B16-OVA screen. **(F)** Identification of top hits enhancing OT1 CD8^+^ T cell infiltration 11 days after transfer into EG7-OVA and B16-OVA tumors using a library of 531 genes encompassing top hits identified in Figure 1B. Except for *Trac*, genes depicted in Figure 1C are statistically significant (raw p value < 0.05) based on Limma-voom analysis. See also Figure S1-2.

Upon bookshelf merging, hits were called based on UMIs (results by bookshelf depicted in Figure S1D). We observed enrichment of sgRNAs targeting positive controls and depletion of negative control sgRNAs (Figure 1B) consistent with our prior screens[31]. Multiple genes were strongly enriched including *Zc3h12a, Ankrd11, Ikzf1, Chic2, Setd5, Prkd2, Peli1, Stub1, Gnas, Rc3h1, Tnfaip3* and *Traf6*. In particular, the *Zc3h12a* gene, encoding the ribonuclease Regnase-1, enriched the strongest across all genes (Figure 1B). The second top hit was *Socs1,* encoding suppressor of cytokine signaling 1 (SOCS1). All hits but one were confirmed to be expressed at TPM > 1 in a murine T cell RNA-Seq dataset that is described later in Figure 3E (Figure S1E). Hit calling based on read counts instead of UMIs yielded similar hits, albeit with a modified hit rank order (Figure S1F).

To define first principles underlying CD8^+^ T cell anti-tumor functionality, pathway commonalities across top hits were annotated (Figure 1C). Regnase-1 and *Rc3h1*, which encodes the RNA binding protein Roquin-1, both direct the degradation or inhibition of mRNA transcripts including negative regulators of NF-κB signaling[37,38], with inactivation in CD8^+^ T cells enhancing anti-tumor function[26,28]. The JAK/STAT and TGFβ cytokine signaling pathways were represented (Figure 1C and Figure S1G) as well as regulators of the TCR signaling pathway, including many known regulators of the NF-κB pathway (Figure 1C and Figure S1G). Transcriptional regulators, including enrichment of *Ikzf1,* which encodes Ikaros and which has been reported to drive a Tex^prog^ state[39], and depletion of *Klf2*, which encodes Klf2 and which maintains Tex lineage fidelity[40] were represented (Figure 1C and S1G). Novel methyltransferases as well as unclassified targets with no previously defined role in CD8^+^ T cells strongly enriched, including *Ankrd11, Gnas,* and *Stk40*. Gene set enrichment analysis indicated IL6/JAK/STAT3 and the TNFα via NF-κB signaling pathways as the two top hallmark pathways (Figure 1D). Genes induced by KRAS activation also were enriched, with *Tnfaip3*, *Trib1*, and *Inhba* as leading-edge KRAS pathway genes showing overlap with TNFα signaling (Figure 1D). Pathways involved in the depletion of CD8^+^ T cells from tumor after removal of DepMap essential genes include oxidative phosphorylation, suggesting a crucial role for this metabolic pathway in sustaining T cell anti-tumor function, consistent with prior screens (Figure 1D)[28].

Additional focused screens were conducted. PMEL[41] TCR-Tg CD8^+^ T cells recognizing gp100 peptide were screened in the MC38-gp100 model, which was previously found to be refractory to PD-1 inhibition[31]. This screen targeted 8,006 genes expressed by T cells and total blood as well as kinases, phosphatases, the ubiquitin pathway, epigenetic as well as metabolic/catabolic proteins. In this screen, *Zc3h12a* and *Socs1* again enriched as the top two hits (Figure 1E, Figure S2A-B, Supplementary Table S2). Lastly, screens were performed in the OT1 / EG7-OVA and OT1 / B16-OVA models using a focused library targeting the top 531 hits identified in the genome-wide screen. In both models, *Zc3h12a* and *Socs1* again scored as the top hits (Figure 1F, Figure S2C, Supplementary Table S3). In conclusion, we systematically mapped the landscape of key pathways and genes regulating CD8^+^ T cell infiltration and accumulation in tumors, identifying the NF-κB and cytokine signaling pathways and Regnase-1 and SOCS1 as the top hits. We additionally confirm the role of the TCR signaling pathway and identify novel proteins not previously implicated in having a functional role in CD8^+^ T cell biology.

### Combination screens identify negative regulators of NF-κB and cytokine signaling as the top dual-edits regulating CD8^+^ T cell function in tumors

To identify dual-edit target combinations enhancing CD8^+^ T cell anti-tumor function beyond single edits, a combination screening approach was used wherein two independent sgRNAs were expressed from the same vector (Figure 2A, Figure S3A). A combination sgRNA library targeting top single-edit hits was constructed (Figure S3B) and evaluated in both the OT1 / B16-OVA and PMEL / MC38-gp100 models (Supplementary Table S4). Comprehensive recovery of sgRNAs from tumor was observed (Figure S3C). Strong enrichment of multiple target pairs was observed, as well as depletion of lethal controls (OT1 / B16-OVA: Figure 2B, PMEL / MC38-gp100: Figure S3D). Of all evaluated combinations, *Rc3h1*, *Zc3h12a, Socs1* and *Nfkbia* were identified as the strongest partners across both models (Figure 2C). We observed target pair interactions wherein enrichment benefits were neutralized similar to a self-pair, such as with *Tgfbr1* with *Tgfbr2*, consistent with both TGFβR1 and TGFβR2 required to transmit TGFβ1 signals (Figure 2D). Traf6 demonstrated neutralizing interactions with *Rc3h1, Peli1,* and *Zc3h12a* and *Tnfaip3*; *Tnfaip3* with *Zc3h12a* and *Peli1*; and *Zc3h12a* with *Nfkbia* (Figure 2E). *Peli1, Traf6, Tnfaip3* and *Nfkbia* encode Pellino-1, Traf6, A20 and IκBα proteins, which are known regulators of NF-κB signaling[42–45]. These neutralizing target-pair interactions suggest that Roquin-1, Pellino-1, Regnase-1, A20, Traf6 and IκBα all regulate CD8^+^ T cell anti-tumor function by in part modulating NF-κB signaling. Indeed, evaluating neutralizing interaction scores as a non-additive interaction network demonstrates clustering of Regnase-1, Roquin and Pellino-1 with NF-κB regulators (Figure 2E).

**Figure 2:**
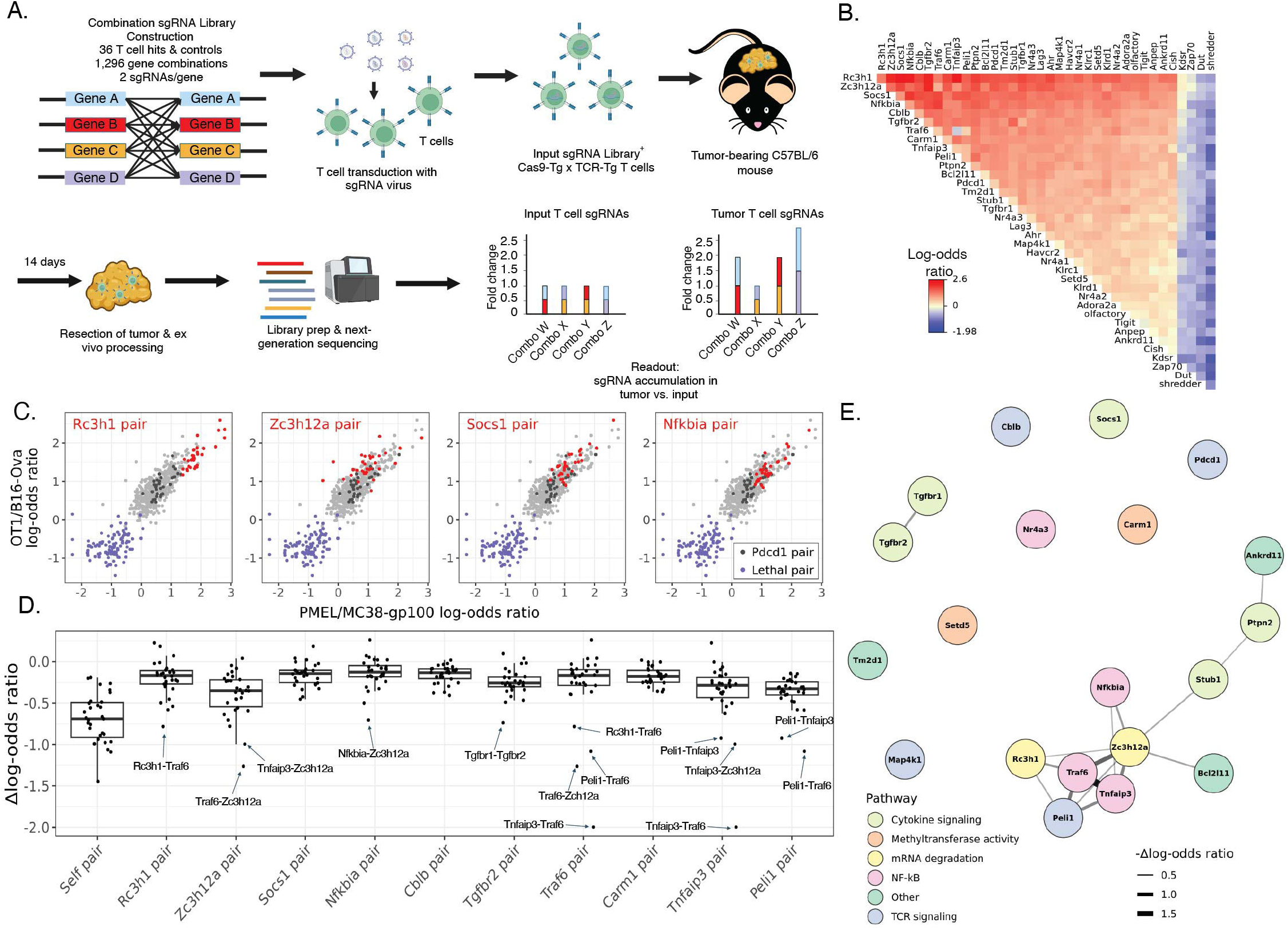
Combination screens identify negative regulators of NF-κB and cytokine signaling as the top dual-edits regulating CD8^+^ T cell function in tumors. **(A)** Experimental schema depicting an in vivo combination CRISPR screen. Viral vectors expressed two sgRNAs targeting top T cell hits combined at random in a pair-wise configuration to generate a combination sgRNA library. sgRNA^+^ TCR-Tg CD8^+^ T cells were transferred into tumor-bearing mice, with tumors harvested 14 days later and the sgRNA distribution of sgRNA^+^ TCR-Tg from tumors and input analyzed. **(B)** Results of a combination screen in the OT1 / B16-OVA model. Target pair enrichment (red) or depletion (blue) of sgRNA pairs are depicted as log-odds ratios in comparison to input using olfactory receptor clone counts as a neutral control. **(C)** Log-odds ratio of OT1 / B16-OVA versus PMEL / MC38-gp100 (shown in Figure S3B) combination screens are depicted for indicated target pairs (red), with *Pdcd1* (dark gray) and lethal (purple) pairs highlighted**. (D)** Combination effects between indicated target pairs were quantified as the difference in the effect size between each target pair in comparison to the sum of the effect sizes for each individual target. Self-pair scores (the scores of each target paired with itself) serve as a control for non-additive effects. **(E)** An interaction network of genes evaluated in combination and scoring at p < 0.05 in the genome-wide screen in Figure 1B, depicting neutralizing interactions (interaction score < 75^th^ percentile of self-pair scores). Edge width represents the absolute value of the interaction score. See also Figure S3.

Candidate pairs were next evaluated in efficacy studies. Given the strong representation of NF-κB and cytokine signaling pathways in the single edit and combination screens, pairs were selected inclusive of representation of both pathways, with Roquin-1, Regnase-1, and IκBα selected to be combined with SOCS1.

### Dual inactivation of SOCS1 and Regnase-1 in transferred CD8^+^ T cells drives tumor regression and prolongs survival in syngeneic tumor models refractory to single edited CD8^+^ T cells

We next evaluated the impact of SOCS1 (sgSocs1) as a dual-edit together with either Regnase-1 (sgRegnase-1), Roquin-1 (sgRoquin-1) or IκBα (sgIκBα) on CD8^+^ T cell anti-tumor function against large, established tumors defined as >300mm^3^ in size and in comparison to single-edits, PD-1 (sgPD-1), and irrelevant controls (sgOlf). All SOCS1 combinations as well as Regnase-1 single edited OT1s demonstrated complete regression of tumor (Figure 3A). Following tumor clearance, mice were re-challenged with B16-OVA and rejected secondary tumor challenge (Figure 3B). SOCS1/IκBα and SOCS1/Roquin-1 dual-edited OT1s presented predominantly as CD44^+^CD62L^+^ T central memory (T_cm_) cells in blood prior to re-challenge and expanded into CD44^+^CD62L^-^ T effector memory (T_em_) cells followed by contraction and continued persistence as CD44^+^CD62L^+^ T_cm_ cells following challenge, similar to prior results with single inactivation of SOCS1 (Figure S4A, D-E)[31]. In contrast, Regnase-1 single and SOCS1/Regnase-1 dual-edited OT1s presented predominantly as T_em_ cells, with minimal expansion observed following re-challenge (Figure S4B-C).

**Figure 3:**
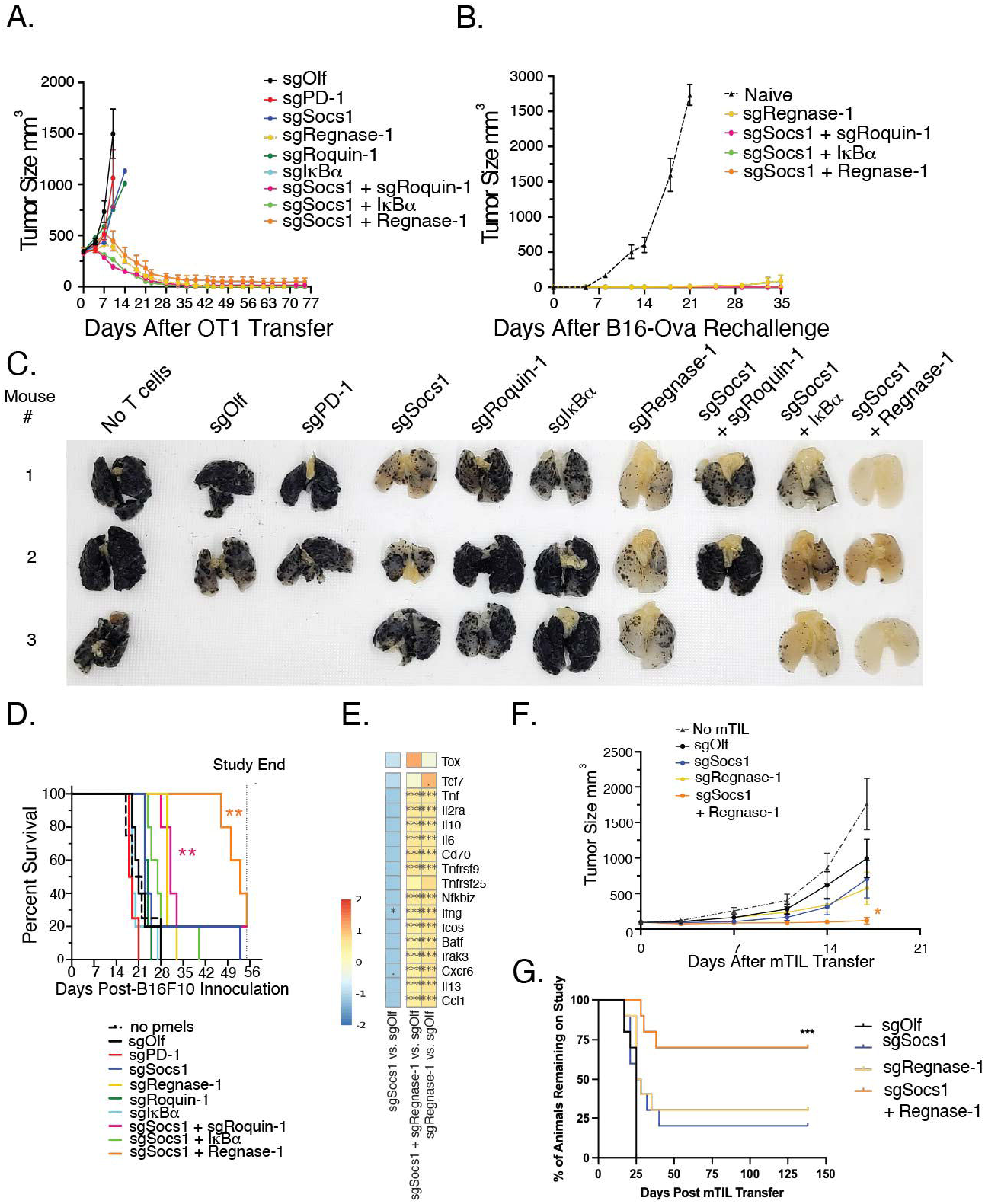
Dual inactivation of SOCS1 and Regnase-1 in transferred CD8^+^ T cells drives tumor regression and prolongs survival in syngeneic tumor models refractory to single edited CD8^+^ T cells. **(A)** C57BL/6 mice bearing ∼350mm^3^ subcutaneous B16-OVA tumors were treated with 3×10^6^ single or dual-edited CD8^+^ OT-1 T cells as indicated. Tumor growth curves of each group over time are depicted. **(B)** Mice undergoing complete tumor rejection in Figure 3A were re-challenged with B16-OVA tumor cells 78 days following initial OT1 transfer with tumor growth by treatment group depicted. **(C)** Mice with disseminated B16F10 tumors were treated with edited PMEL CD8^+^ T cells as indicated. Lungs were harvested on Day 15 from 2-3 mice as represented and evaluated for tumor burden. **(D)** Same experimental design as in Figure 3C, with n=5 mice/group evaluated for survival benefit over time. **(E)** Selected top DEGs between single and dual-edited mTIL derived from B16-OVA tumors were determined by RNA-Seq, with pairwise log fold changes depicted by heat-map. **(F)** Single or dual-edited mTIL were adoptively transferred into C57BL/6 mice bearing B16-OVA tumors, with tumor growth curves depicted by treatment group depicted. **(G)** Overall survival benefit by mTIL treatment group. Statistical significance between treatment groups compared to sgOlf determined using a Mantel-Cox test in Figure 3D and H and a two-way ANOVA in Figure 3G, with * = p value < 0.05 and ** = p value < 0.01. For Figure 3E, the *s in the heatmap represent the adj-p value (Benjamini Hochberg) for pairwise differential expression analysis with DESeq2 where *** < 0.001 and * 0.01-0.05. Figure 3A is representative of n=2, Figure 3C is representative of n=2, and Figure 3G-H are representative of n=2 similar studies. See also Figure S4-5.

The ability of single and dual-edited PMELs to prolong survival in a disseminated B16F10 model was evaluated. SOCS1/Regnase-1 dual-edited PMELs led to reduction in lung tumor burden (Figure 3C) and increased survival benefit in comparison to SOCS1/Roquin-1 and SOCS1/IκBα (Figure 3D). Regnase-1 single edits additionally reduced tumor burden (Figure 3C) and improved survival (Figure 3D), while SOCS1, Roquin-1, PD-1, IκΒα and Olf single edited PMELs provided no discernable benefit (Figure 3C-D).

To evaluate the impact of inactivating SOCS1 and Regnase-1 in Tumor Infiltrating Lymphocytes (TIL), we developed a murine TIL model (mTIL). mTIL isolated from B16-OVA tumors (Figure S5A) were CD8^+^ and displayed reactivity to B16-OVA with ∼6% of cells staining for SIINFEKL tetramer (Supplemental Figure S5B-C). RNA-Seq analysis displayed enhanced Regnase-1-driven expression of *Il2ra, Tnfrsf9*, *Irak3* and *Ifng* as well as Regnase-1 substrates *Icos, Nfkbiz,* and *Batf*[28,33,46] in both single and dual-edited mTIL (Figure 3E). Heightened expression of 4-1BB, CD25 and ICOS by the Regnase-1 single and dual-edited mTIL was confirmed by flow cytometry (Figure S5D). Following transfer into B16-OVA tumor-bearing mice, impaired tumor growth was observed with Regnase-1 and SOCS1 single edits, with dual-edited mTIL displaying marked enhancement in long-term survival and anti-tumor activity (Figure 3F-G; individual mice in Figure S5E). Body weights of all mice with complete tumor regressions were monitored, with no loss observed (Figure S5F).

By evaluating the top combinations representing the NF-κB and cytokine signaling pathways in TCR-Tg and mTIL models, we demonstrate that Regnase-1 inactivation drives the strongest CD8^+^ T cell anti-tumor activity amongst evaluated single edits and that SOCS1/Regnase-1 dual-edits drives the strongest efficacy amongst evaluated dual-edits. Inactivation of Regnase-1 in a single and dual-edit setting additionally gave rise to persistent and functional T_em_ cells.

### SOCS1 and Regnase-1 inactivation cooperate to expand Tex^int^ and Tex^eff^ cells in lymphoid tissues and tumor and form persistent T_em_ cells following tumor clearance

We next investigated how SOCS1 and Regnase-1 inactivation impact CD8^+^ T cell subsets during the acute and memory phases of the anti-tumor response. We transferred PD-1, SOCS1, and Regnase-1 single and SOCS1/Regnase-1 dual-edited OT1s into mice bearing >350mm^3^ B16-OVA tumors, with SOCS1 inactivation delaying tumor growth and Regnase-1 single and SOCS1/Regnase-1 dual-edits driving complete tumor regression (Figure S6A-C). The rapid tumor regression observed impeded ex vivo characterization of OT1s (Figure S6A), therefore, we characterized single and dual-edited OT1s from mice bearing tumors ∼560mm^3^ in size on the day of transfer. Regnase-1 single edits enhanced OT1 accumulation in blood and spleen with low levels of accumulation observed by all other single edits and controls (Figure 4A). Relative to Regnase-1 single edits, dual-inactivation of Regnase-1 and SOCS1 more than doubled OT1 accumulation in blood and spleen (Figure 4A). In tumor draining lymph nodes (TDLN), Regnase-1 single and dual-edited OT1s accumulated similarly, with little accumulation again observed by other single edits and controls. Within tumor, and consistent with our combination screen results, dual-edited OT1s comprised most of CD8^+^ T cells (Figure 4A). These data demonstrate that SOCS1 and Regnase-1 cooperate to enhance the accumulation of tumor-specific CD8^+^ T cells in lymphoid tissues and tumor during the acute phase of the anti-tumor response.

**Figure 4:**
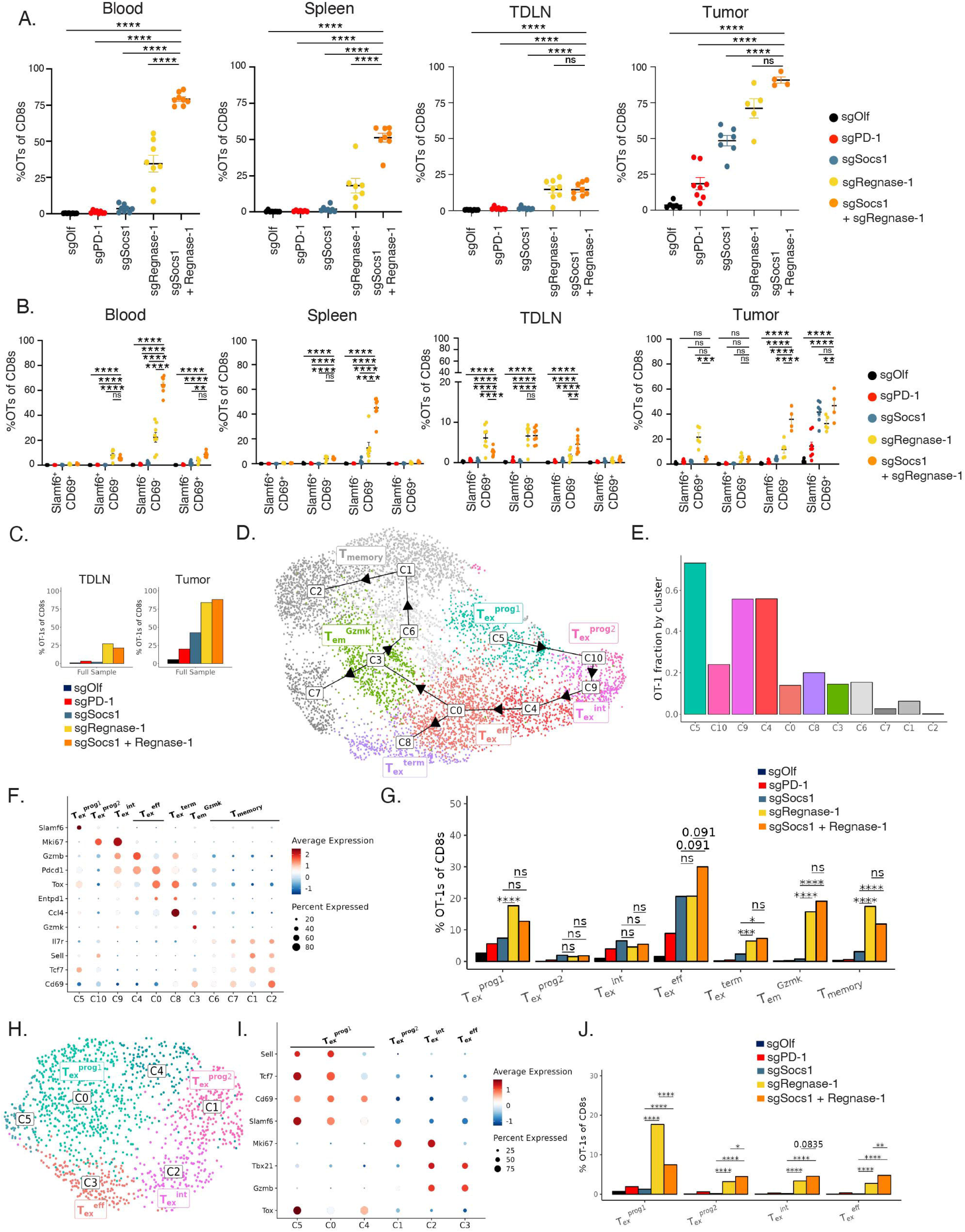
SOCS1 and Regnase-1 inactivation cooperate to expand Tex^int^ and Tex^eff^ cells in lymphoid tissues and tumor. CD45.1^+^ mice bearing 560mm^3^ B16-OVA tumors were treated with 3×10^6^ single or dual-edited CD8^+^ OT-1 T cells as indicated. **(A)** OT1 frequency in blood, spleen, tumor draining lymph nodes (TDLNs) or tumor 7 days following transfer. **(B)** Slamf6^+^CD69^+^, Slamf6^+^CD69^-^, Slamf6^-^CD69^-^ and Slamf6^-^CD69^+^ OT1s as a frequency of CD8^+^ T cells was quantified from the indicated tissue between treatment groups. **(C)** Frequency of OT1s of total CD8^+^ T cells by single or dual edits quantified by 5’ scRNA-Seq of CD45^+^ cells isolated from tumor or TDLNs, as indicated. **(D)** UMAP visualization of tumor CD8^+^ cells by cluster, with Slingshot analysis depicting differentiation trajectories. Cluster annotations are based on expression of Tex subset-defining signatures. **(E)** Tumor OT1s as a frequency of CD8^+^ T cells by cluster, with cluster order determined by differentiation trajectory. **(F)** Gene expression Dot plot by indicated cluster in tumor. **(G)** Frequency of tumor OT1s of total CD8^+^ T cells by cluster and by treatment group. **(H)** UMAP visualization of TDLN OT1 cells, with clusters annotated based on Tex subset-defining signatures. **(I)** Gene expression Dot plot by indicated cluster in TDLNs. **(J)** Frequency of TDLN OT1s of total CD8^+^ T cells by cluster and by treatment group. Statistical significance between treatment groups determined using a two-way ANOVA in Figures 4A and 4B and using a chi-squared test with Holm correction in Figures 4G and 4J, with ns = no significance, * = p value < 0.05, ** = p value < 0.01, *** = p value < 0.001 and **** = p value < 0.0001. Experiment depicted in Figure 4A-B representative of n=2 similar studies. See also Figure S6-8.

To evaluate the impact of single and dual edits on Tex subsets, we adapted an approach based on Slamf6/CD69 expression as described by Beltra et al[13]. Slamf6 identifies Tcf1-expressing Tex^prog^ cells[13,47], with quiescent and resident progenitor Tex^prog1^ cells identified by Slamf6^+^CD69^+^, proliferating and circulating progenitor Tex^prog2^ cells identified by Slamf6^+^CD69^-^, proliferating and circulating Tex^int^ cells identified by Slamf6^-^CD69^-^ and terminally exhausted and resident Tex^term^ cells identified by Slamf6^-^CD69^+^. Consistent with Regnase-1 directly targeting *Tcf7* transcripts[48], Regnase-1 single and SOCS1/Regnase-1 dual-edit OT1s increased the accumulation of Slamf6^+^CD69^+^ Tex^prog1^ cells in TDLN and Slamf6^+^CD69^-^ Tex^prog2^ cells in all tissues (Figure 4B). Interestingly, dual-inactivation of Regnase-1 and SOCS1 drove less of an increase of Tex^prog1^ cells in TDLN and with no Tex^prog1^ cells observed in tumor compared with single inactivation of Regnase-1 (Figure 4B). Inactivation of Regnase-1 enhanced Slamf6^-^CD69^-^ Tex^int^ cells, while SOCS1/Regnase-1 dual-edits strongly enhanced accumulation of this subset in blood, spleen, TDLNs and tumor (Figure 4B). Slamf6^-^CD69^+^ Tex^term^ subsets were primarily detected in tumor (Figure 4B), where SOCS1 and Regnase-1 single edits and SOCS1/Regnase-1 dual-edits all enhanced accumulation of this subset to a similar degree (Figure 4B). Using CD44 and CD62L as additional phenotypic markers, Regnase-1 single and, to a lesser degree, dual-edits enhanced the accumulation of CD62L^+^CD44^+^ T_cm_ cells in TDLN (Figure S6D) while dual edits strongly enhanced the accumulation of CD62L^-^CD44^+^ T_em_ cells in all tissues analyzed. Based on these collective flow cytometry data, we conclude that SOCS1 and Regnase-1 dual-edits enhance the accumulation of Slamf6^-^CD69^-^Tex^int^ cells in blood, spleen, TDLN and tumor.

To further elucidate how SOCS1 and Regnase-1 inactivation impacts Tex subsets, paired single-cell 5’ scRNA-Seq/scTCR-Seq was performed on CD45^+^ lymphocytes from tumor and TDLN. From tumor, 54,554 independent transcriptomes were annotated (Figure S7A) for immune cell lineages based on transcript expression by cluster (Figure S7B). Analysis of the *Cd3d^+^* cluster (Figure S7C) revealed *Cd8a*, *Cd4,* and cells expressing *Foxp3* (Figure S7D). Quantifying Treg frequency revealed a notable reduction by Regnase-1 single and SOCS1/Regnase-1 dual edits (Figure S7E). The frequencies of OT1 cells as a fraction of CD8^+^ T cells in TDLNs and tumor were consistent with our flow cytometry analysis (Figure 4C), with Regnase-1 single and SOCS1/Regnase-1 dual edits both showing the strongest enrichment (Figure 4C). Tex states occupied by 7,005 tumor-localized CD8 transcriptomes were mapped, with Slingshot[49] used to determine differentiation trajectory across clusters (Figure 4D). OT1 cells were detected in all clusters, comprising more than 50% of CD8^+^ T cells in clusters C5, C9 and C4 (Figure 4E). We annotated clusters based on Tex-defining lineage transcripts (Figure 4F, Figure S7F-G) as well as by comparing cluster gene expression signatures with gene sets used by Beltra et al. and Miller et al. (Figure S7H-I)[13,47]. OT1-enriched clusters were mapped to a differentiation trajectory comprising Tex^prog1^, Tex^prog2^, Tex^int^, Tex^eff^, and Tex^term^ (Figure 4D, F). We and others have described *Mki67^-^Gzmb^+^*effector Tex^eff^ cells marked by intermediate expression levels of *TOX*[31,47]. Our Tex subset annotation confirms a Slamf6^-^CD69^-^ phenotype encompasses Tex^int^ and Tex^eff^ cells, and a Slamf6^-^CD69^+^ phenotype includes Tex^term^ cells (Figure 4F). Clusters enriched for endogenous CD8^+^ T cells (C6, C7, C1 and C2; Figure 4D-E) contained Gzmk^+^ effector memory (Tem^Gzmk^) and memory (T_memory_) cells expressing *Il7r*. These subsets lacked *Slamf6* but expressed *Cd69* (Figure 4F), indicating that Slamf6^-^CD69^+^ OT1s in tumor encompasses Tem^Gzmk^ and T_memory_ cells in addition to Tex^term^ cells.

We evaluated how single and dual-inactivation of SOCS1 and Regnase-1 impacted the accumulation of each subset. Regnase-1 inactivation broadly enhanced the accumulation of all Tex states as well as uniquely expanded Tem^gzmk^ and T_memory_ cells. Inactivation of SOCS1 drove the accumulation of Tex^int^ and Tex^eff^ cells, consistent with our prior work[31]. Relative to Regnase-1 single edits, SOCS1/Regnase-1 dual-edits further increased Tex^eff^ cells within tumor (Figure 4G).

In TDLNs, we clustered OT1s both with endogenous CD8^+^ T cells (Figure S7K) as well as without (Figure 4H). Our analyses focused on co-clustered OT1s (Figure 4H) due to the high frequency of different transcriptional states occupied by endogenous CD8^+^ T cells. Tex subsets were again identified based on marker genes (Figure 4I and Figure S7L). Regnase-1 single edits again broadly increased Tex subset accumulation, and particularly Tex^prog1^ cells. Like in tumor, SOCS1/Regnase-1 dual-edits again further increased the accumulation of Tex^eff^ subsets in TDLN while displaying diminished enhancement of Tex^prog1^ cells versus Regnase-1 single edit (Figure 4J). Extending our flow cytometry analysis which found dual inactivation of SOCS1/Regnase-1 enhancing the accumulation of Slamf6^-^CD69^-^ Tex^int^ cells in blood, spleen, TDLN and tumor, these results demonstrate that Slamf6^-^CD69^-^ Tex^eff^ cells are additionally increased in TDLN and tumor.

Finally, we confirmed the impact of SOCS1 and Regnase-1 single and dual-edits on the memory phase of the CD8^+^ T cell anti-tumor response. Complete responder mice from Figure S6A-C were rechallenged with B16-OVA and again rejected tumor (Figure S8A). SOCS1 edited OT1s in the peripheral blood again exhibited a T_cm_ phenotype, expanded into T_em_ cells following re-challenge, and contracted to regain a T_cm_ phenotype (Figure S8B-D). Regnase-1 single edited OT1s were detected at lower frequencies and with a heterogenous T_cm_/T_em_ phenotype prior to and following re-challenge. SOCS1/Regnase-1 dual-edited cells were detected at a higher baseline frequency and with a relatively homogenous T_em_ phenotype in comparison to the Regnase-1 single edits. Following tumor rechallenge, dual-edited OT1s continued to persist at steady-state predominantly as T_em_ cells (Figure S8C-D). These data demonstrate that SOCS1/Regnase-1 dual-edits give rise to persistent T_em_ memory cells at increased frequency in comparison to Regnase-1 single edit following tumor clearance.

These results demonstrate that in the acute phase of the anti-tumor response, dual-inactivation of SOCS1 and Regnase-1 cooperate to strongly increase the accumulation of Slamf6^-^CD69^-^ Tex^int^ cells in the blood and spleen and Tex^eff^ cells within TDLN and tumor. In the memory phase and following tumor clearance, dual inactivation of SOCS1 and Regnase-1 facilitated the formation of persistent T_em_ cells.

### Dual-inactivation of SOCS1 and Regnase-1 rewires Tex subsets through non-overlapping mechanisms to promote an effector state while reducing regulators of exhaustion

We next determined the impact of SOCS1 and Regnase-1 inactivation on CD8^+^ Tex subsets by characterizing their transcriptional state. Using flow cytometry, SOCS1/Regnase-1 dual-edited OT1s were found to express high levels of T-bet^+^ and Ki67^+^ in blood, spleen, and TDLN in comparison to single edits (Figure 5A, S9A). T-bet is a key mediator of effector function in CD8^+^ T cells whose expression together with Ki67 is a hallmark of Tex^int^ cells[13], with these data confirming Slamf6^-^CD69^-^ OT1s in blood and spleen (Figure 4B) as Tex^int^ cells as well as a majority of OT1s within TDLN, with the magnitude of Tex^int^ cells in TLDN not fully captured in the scRNA-Seq data. Regnase-1 single and dual-edits additionally drove heightened expression of Icos, with *Icos* transcripts substrates of Regnase-1 ribonuclease activity[33]. Single inactivation of Regnase-1 was unique in enhancing Tcf1 expression in TDLN OT1s, consistent with Regnase-1 single edits enhancing accumulation of Tex^prog^ cells (Figure 4B). Of interest, dual-edited OT1s exhibited reduced expression of the exhaustion regulators Eomes[50,51] and TOX[52–55] within TDLN OT1s (Figure 5A).

**Figure 5:**
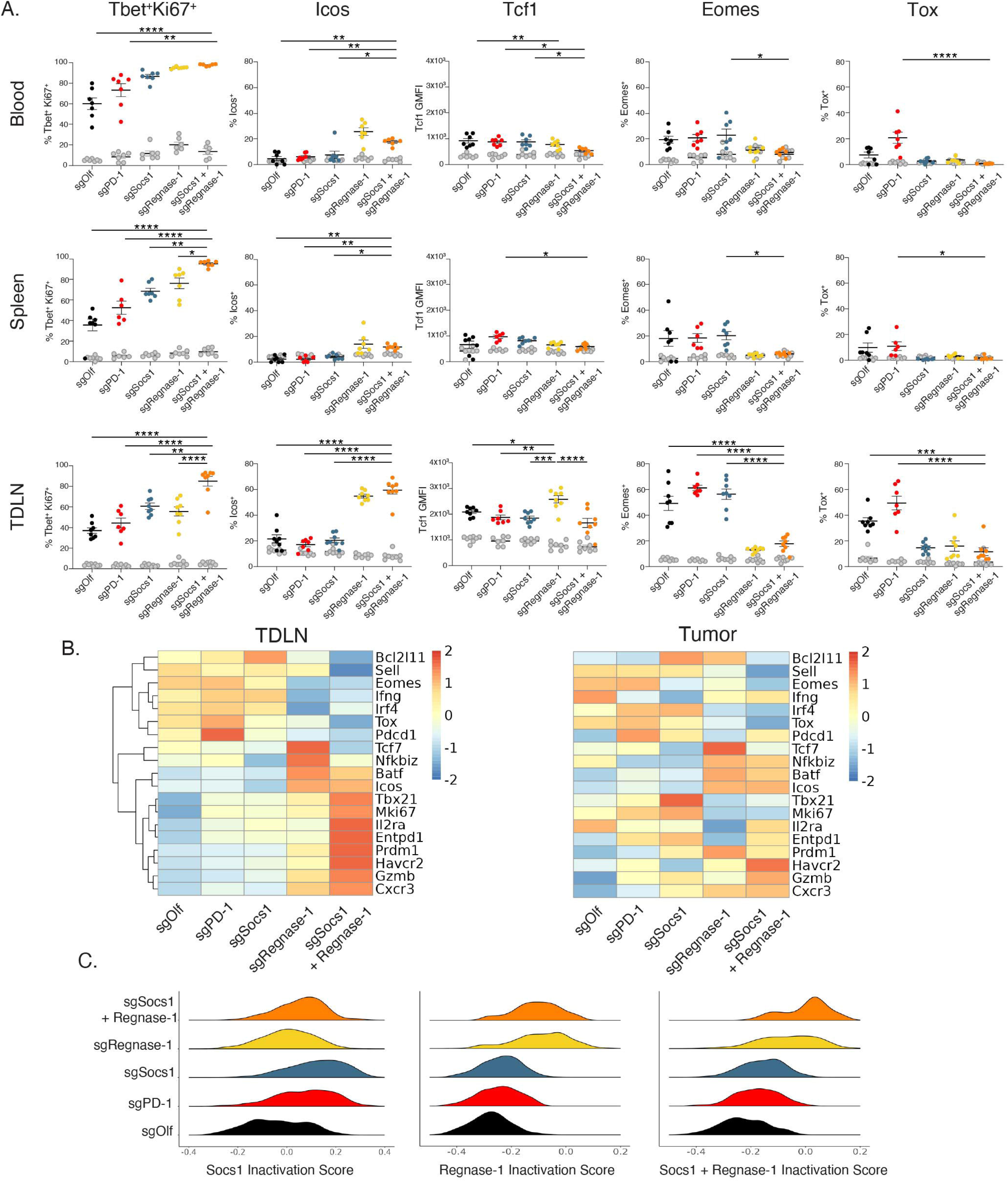
Dual-inactivation of SOCS1 and Regnase-1 rewires Tex subsets through non-overlapping mechanisms to promote an effector state while reducing regulators of exhaustion. SOCS1 and Regnase-1 single and dual-edited OT1s were characterized seven days following transfer into B16-OVA tumor-bearing mice. (A) Expression of the indicated protein by OT1s from the indicated tissue by treatment group using flow cytometry. Colored data points indicate gated OT1s, gray data points indicate gated endogenous CD8^+^ T cells. (B) Pseudo-bulk RNA-Seq analysis of single and dual-edited TDLN and tumor-derived OT1s depicting expression of the indicated transcripts. (C) SOCS1, Regnase-1, and SOCS1 + Regnase-1 inactivation gene sets were used to score single and dual-edited OT1s as indicated. Statistical significance between treatment groups determined using a one-way ANOVA in Figures 5A, with * = p value < 0.05, ** = p value < 0.01, *** = p value < 0.001 and **** = p value < 0.0001. See also Figure S9.

Pseudo-bulk RNA-Seq of TDLN and tumor OT1s demonstrated that dual-inactivation of SOCS1/Regnase-1 strongly enhanced the expression of transcripts associated with effector function including *Il2ra, Havcr2, Gzmb* and *Cxcr3* (Figure 5B). As observed in our flow cytometry data, dual edits increased *Mki67* and *Tbx21* expression in TDLN OT1s while tumor OT1s conversely displayed reduced expression. Regnase-1 single and dual edits exhibited decreased *Ifng* in TDLN OT1s, and conversely increased expression in tumor. Regnase-1 single and dual-edits enhanced the expression of Regnase-1 substrates *Nfkbiz, Batf* and, confirming our flow cytometry data, *Icos*. The exhaustion regulators *Irf4*[56] and, consistent with our flow cytometry data, *TOX* and *Eomes* were downregulated in the SOCS1/Regnase-1 dual-edit TDLN and tumor-derived OT1s (Figure 5B). Lastly, SOCS1 and Regnase-1 dual inactivation suppressed the expression of *Bcl2l11*, which encodes BIM which functions to cull the CD8^+^ T effector cell pool through apoptosis[57], in comparison to single edits.

These data indicate substantial changes in the transcriptional state of Tex cells at the population level. We additionally explored changes at the Tex subset level using scRNA-Seq (Figure S9B). Within tumor, we were limited in our ability to evaluate Tex^prog2^ subsets as well as Olf and PD-1-edited Tex^term^, Tex^Gzmk^ and T_memory_ subsets given low cell counts. Therefore, the few Tex^prog2^ cells were combined with Tex^int^ cells into a ‘proliferating’ (Tex^prolif^) subset, with Olf and PD-1 groups omitted from the Tex^term^, Tex^Gzmk^ and T_memory_ subset comparisons (Figure S9B). We observed an impact of Regnase-1 single and dual edits even within discrete Tex subsets. Dual edits increased *Tcf7* and *Slamf6* in Tex^prog1^ cells, and additionally increased *Batf, Il2rg, Prf1* and *Gzmb* while reducing *Tcf7, Slamf6, Irf4* and *Bcl2l11* within Tex^int^/Tex^prolif^ and Tex^eff^ subsets in TDLN and tumor OT1s. *Bcl2l11* expression was additionally reduced in T_memory_ cells. Heightened expression of *Tbx21* was observed in TDLN Tex^int^ and Tex^eff^ subsets, and decreased expression within Tex^prolif^ and Tex^eff^ subsets within tumor. Within Tex^term^ cells in tumor, Regnase-1 single and dual-edits increased the expression of *Il2rg*, *Nfkbiz, Cxcr3, Ifng* and *Gzmk*, as well as *Gzmk* and *Cxcr3* within Tem^Gzmk^ cells. Regnase-1 single and, in particular, dual-inactivation reduced *Eomes* and *Tox* expression across Tex subsets, including the Tex^term^ subset, which typically expresses high TOX. We confirmed Tex subset-specific increases of CD25 and TIM3 by SOCS1/Regnase-1 dual edits by flow cytometry (Figure S9C).

Pairwise comparison of gene expression signatures between single and dual-edited OT1s from TDLNs and tumor was performed. Evaluating DEG (differentially expressed genes) overlap between either single and dual-edit treatment (Supplementary Tables S5-S7) groups versus Olf revealed little overlap between single edit versus Olf comparisons (Figure S9D), indicating independent contributions by SOCS1 and Regnase-1 to dual-edit functionality. Greater overlapping DEGs were observed between Regnase-1 single and SOCS1/Regnase-1 dual edits when compared to SOCS1 single and SOCS1/Regnase-1 dual edits (Figure S9D), indicating Regnase-1 impacts CD8^+^ T cell function greater than SOCS1. To additionally evaluate the contributions of SOCS and Regnase-1 gene signatures to dual-edit functionality, we generated SOCS1, Regnase-1 and SOCS1/Regnase-1 inactivation gene sets (Supplementary Tables S8-10) and scored single and dual-edited OT1s (Figure 5C). The distributions of scores reveal mechanistic overlap between PD1 and SOCS1, consistent with our prior work[31], and demonstrate that SOCS1 and Regnase-1 inactivation are non-overlapping in their functional impact on CD8^+^ T cells.

These data demonstrate that dual inactivation of SOCS1 and Regnase-1 rewires Tex subsets by enhancing the expression of effector regulators in lymphoid tissue and tumor-derived Tex and memory subsets, and by reducing the expression of transcriptional regulators driving terminal exhaustion. Lastly, we demonstrate that SOCS1 and Regnase-1 impact the functional state of CD8^+^ T cells through cooperative and non-overlapping mechanisms.

### Inactivation of SOCS1 and Regnase-1 enhances the anti-tumor functionality of human TIL and CAR-T cell therapies

We extended our findings on the strong anti-tumor activity observed by SOCS1/Regnase-1 dual-edits in murine CD8^+^ T cells to human Tumor Infiltrating Lymphocyte (TIL) and CAR-T therapies in solid tumor models. In tumor samples obtained from either naïve or pre-treated patients with advanced metastatic melanoma (n=3), HNSCC (n=1) and NSCLC (n=3) (Supplementary Table S11), CRISPR/Cas9 engineered eTIL were manufactured including inactivation of SOCS1 in KSQ-001EX, a previously described experimental therapy[31], Regnase-1 single edits, and dual-inactivation of SOCS1 and Regnase-1 in KSQ-004EX. No Electroporation (No EP) controls were additionally included. The cellular characterization of KSQ-001EX and KSQ-004EX eTIL was CD3^+^ (Figure S10A), CD8^+^ with few CD4^+^ cells (Figure S10B-C) and displaying a CD45R0^+^CCR7^-^ T effector memory phenotype (Figure S10D) with nearly complete inactivation of the *SOCS1* and *ZC3H12A* genes (Figure S10E). Upon co-culture with A375-mOKT3 tumor spheroids[31], KSQ-001EX and KSQ-004EX both displayed improved abilities to kill and produce IFNγ (Figure 6A-B). These data suggest that SOCS1 inactivation is the primary driver enhancing these functions. eTIL manufactured from a checkpoint inhibitor-pretreated melanoma donor were repeatedly stimulated by spheroids comprised of A375 cells engineered to express membrane-associated OKT3 (A375-mOKT3). Both KSQ-001EX and Regnase-1 single edit displayed an increased ability to kill and with KSQ-004EX displaying an even more pronounced ability to kill, indicating that inactivation of SOCS1 and Regnase-1 combine to enhance KSQ-004EX refractoriness to chronic stimulation (Figure 6C).

**Figure 6:**
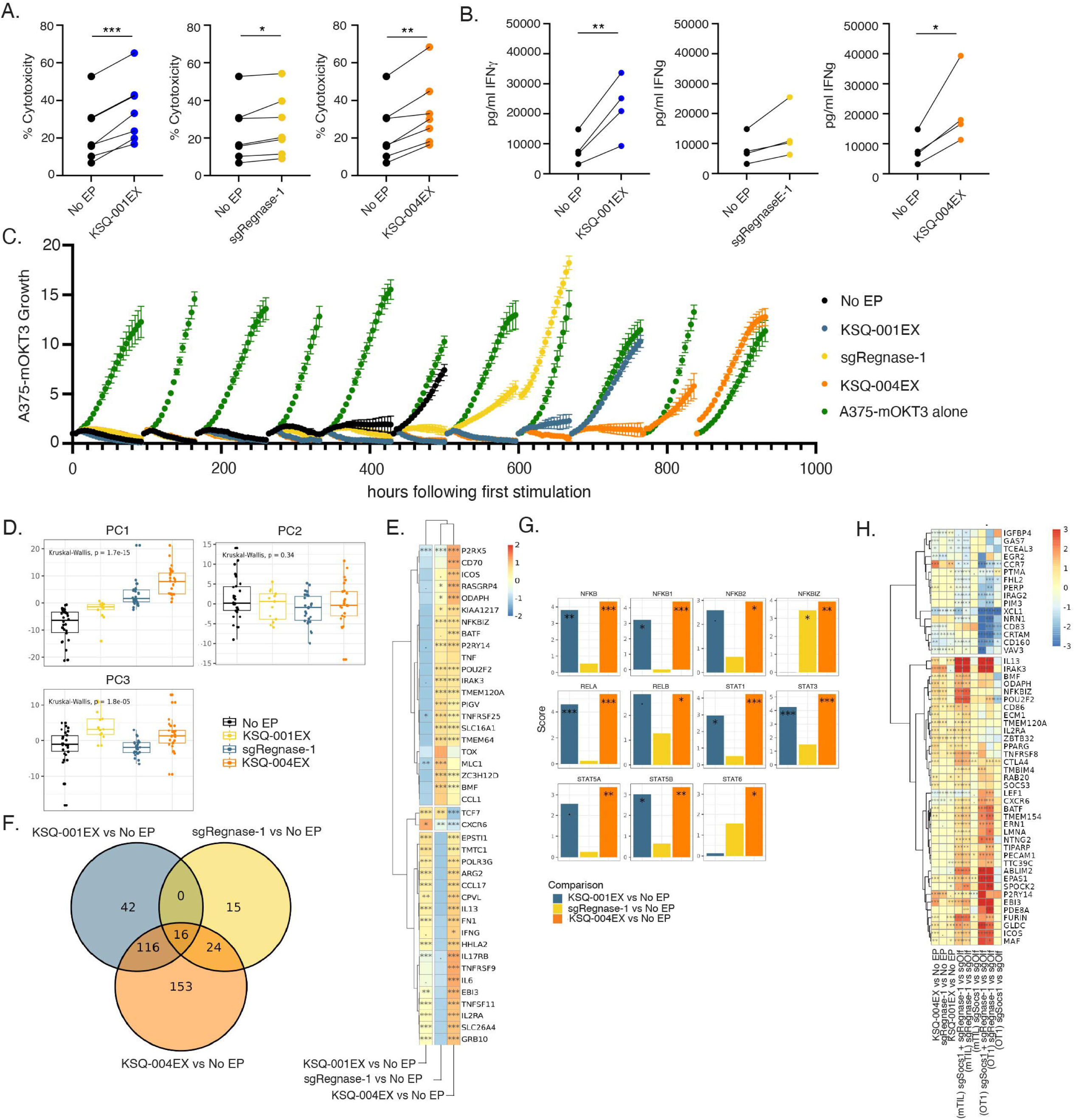
Inactivation of SOCS1 and Regnase-1 enhances the anti-tumor functionality of human TIL. CRISPR/Cas9 engineered TIL (eTIL^®^) were manufactured to inactivate SOCS1 (KSQ-001EX), Regnase-1 (sgRegnase-1), or both SOCS1 and Regnase-1 (KSQ-004EX) with non-electroporated controls used as a benchmark (No EP). **(A)** Percent killing of A375-mOKT3 tumor spheroids by indicated eTIL. **(B)** Production of IFNγ from during eTIL co-culture with A375-mOKT3 tumor spheroids. **(C)** Repeat stimulation assay assessing the ability of eTIL manufactured from a treatment-refractory melanoma donor to kill A375-mOKT3 cells following each stimulation over time. **(D)** eTIL bulk RNA-Seq, with Principle Components (PC) PC1, PC2 and PC3 depicted. No EP contains 30 independent donors; KSQ-001EX contains 26 independent donors; sgRegnase-1 contains 13 independent donors, and KSQ-004EX contains 25 independent donors. **(E)** Select DEGs between pairwise eTIL comparisons are depicted, with log fold change depicted by heatmap. **(F)** The number of DEGs between the eTIL comparisons are depicted by Venn diagram. **(G)** Transcription factor activity analysis from eTIL bulk RNA-Seq **(H)** Heatmap of log fold changes of top DEGs between edited TILs versus their respective controls in both human and mouse. Statistical significance between treatment groups was determined using a two-way ANOVA in Figures 6A-B with ns = no significance, * = p value < 0.05, ** = p value < 0.01, and *** = p value < 0.001. For Figure 6E and 6H, the * in the heatmap represent the adj-p value (Benjamini Hochberg) for pairwise differential expression analysis with DESeq2 where *** = adj-p value < 0.001; ** = 0.001- 0.01; * = 0.01-0.05 and ‘.’ = 0.05-0.1. For Figures 6G, the *s represent adj-p values (Benjamini Hochberg) for transcription factor activity analysis. Figure 6A-B is representative of n=16 independent donors, Figure 6C is representative of n=7 independent donors. See also Figure S10.

We performed bulk RNA-Seq on eTIL, with distinct individual impacts of SOCS1 and Regnase-1 inactivation observed on the transcriptional states of KSQ-001EX and Regnase-1 single edits, and with KSQ-004EX manifesting a combination of both SOCS1 and Regnase-1 biology (Figure 6D). In contrast to murine dual-edited CD8^+^ T cells (Figure 3E, S9D), where Regnase-1 inactivation had a stronger impact on the OT1 transcriptional state, SOCS1 inactivation had a stronger impact on KSQ-004EX. Indeed, greater DEGs were observed between KSQ-001EX versus No EP when compared to sgRegnase-1 versus No EP comparisons, with a significant portion of KSQ-001EX versus No EP DEGS shared with KSQ-004 versus No EP (Figure 6F). Top DEGs include the Regnase-1 substrates *ICOS, NFKBIZ,* and *BATF* as well as *IRAK3*, which were additionally observed in mTIL and OT1s, with SOCS1 driving expression of *CCL17, IL-13, IFNG* and *IL2RA* (Figure 6E). Transcription factor activity analysis (Figure 6G) shows SOCS1-driven enhancement of NF-κB and STAT transcription factor activity, and with Regnase-1 driving increased of the transcriptional regulator NFKBIZ. Analysis of top DEGs across single and dual-edited murine TIL (from Figure 3E) OT1 (from Figures 4-5) and human TIL (from Figure 6E) versus controls (Figure 6H) show commonality of gene signatures across T cell types and species including increased expression of *NFKBIZ*, *IRAK3*, *ICOS* transcripts in Regnase-1 and dual-edited cells and increased cytokine signaling in the SOCS1 and dual-edited T cells.

We next evaluated the anti-tumor activity of single and dual-edited CRISPR/Cas9-edited CAR-Ts (eCAR-Ts) targeting mesothelin (meso eCAR-T) using the M5 CAR[58,59] in the HCT-116 colorectal tumor model. As SOCS1 is a negative regulator of cytokine signals, we sought to appropriately model the availability of cytokines in these studies to be similar to those reported in the clinic. Patients receiving CAR-T therapies typically undergo lymphodepletion resulting in increased cytokine availability, including IL-15[60]. We found that the NSG-Tg(Hu-IL15) strain from Jackson Laboratory expressed a mean of 30pg/ml of IL-15 in blood (Figure S11A), which is similar to amounts present in patients following lymphodepletion at the time of CD19-targeting CAR-T infusion[61]. Meso eCAR-Ts displayed high editing efficiency and CAR expression (Figure S11B), with SOCS1 and Regnase-1 single and dual-inactivation maintaining a balanced CD4/CD8 ratio (Figure S11C) and retaining a Tcm cell phenotype (Figure 7A). Single and dual-edits both enhanced the cytotoxicity of meso eCAR-Ts upon co-cultured with HCT-116 target cells in vitro (Figure 7B).

**Figure 7:**
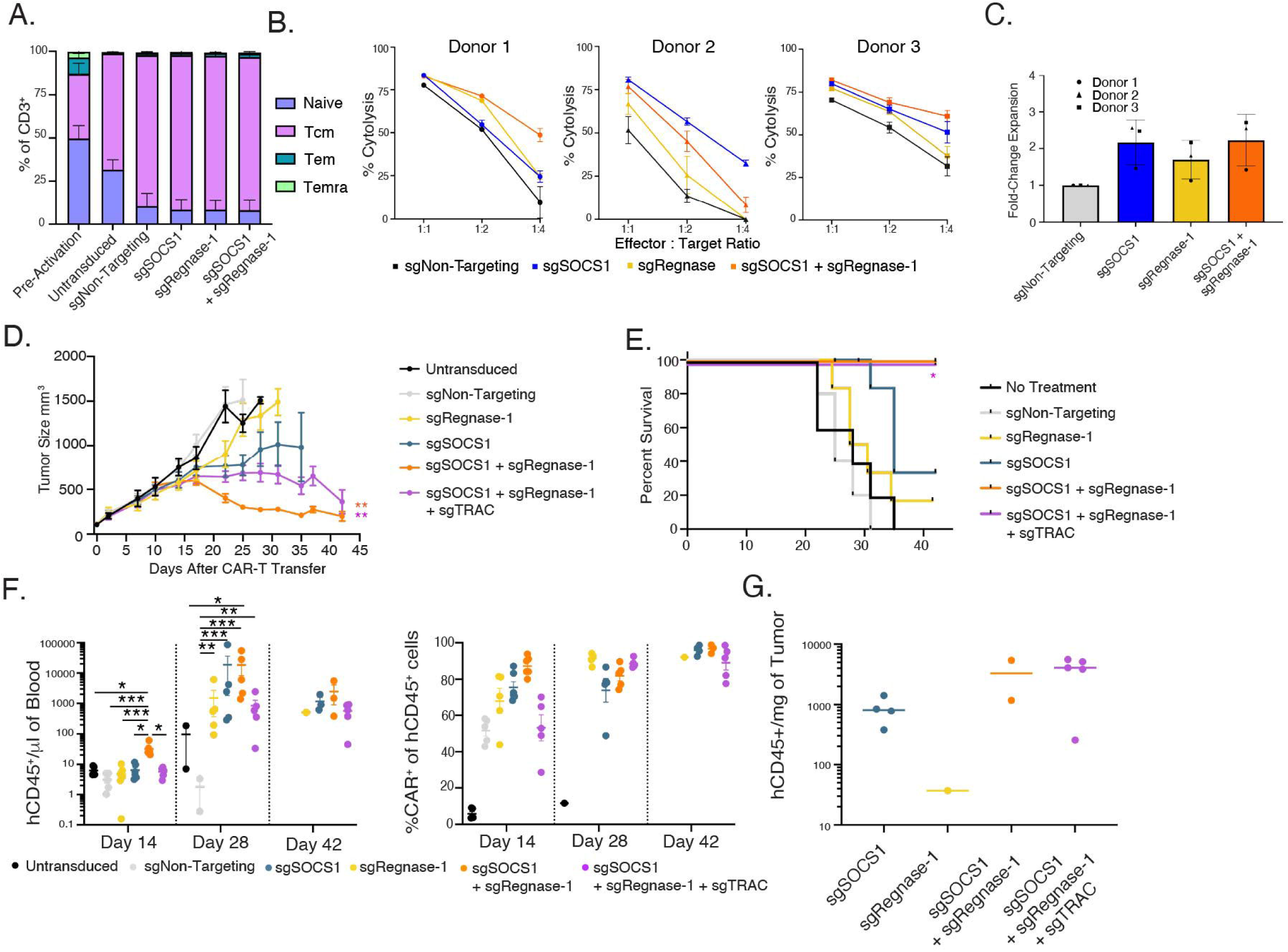
Inactivation of SOCS1 and Regnase-1 enhances the anti-tumor functionality of human meso CAR-Ts. Meso eCAR^™^-Ts were manufactured from n=3 donors and characterized. **(A)** Frequency of naïve (CCR7^+^CD45RA^+^), Tcm (CCR7^+^CD45RA^-^), Tem (CCR7^-^CD45RA^-^), and Temra (CCR7^-^CD45RA^+^) subsets in single and dual-edited meso eCAR-Ts as indicated across donors. **(B)** Percent killing of HCT-116 cells by indicated meso eCAR-Ts. **(C)** Expansion of indicated meso eCAR-Ts in response to HCT-116 target cells. **(D)** Meso eCAR-Ts were transferred into NSG-HuIL-15 Tg mice bearing HCT-116 tumors, with tumor volume shown over time. **(E)** Tumor-burden based survival of Figure 7D. **(F)** Levels of meso eCAR-Ts by treatment group in the peripheral blood on the indicated timepoints (left graph), as well as the proportion of CAR-expressing human CD45+ cells (right graph). **(G)** Absolute number of hCD45^+^ cells in tumor on Day 42 following transfer. For Figure 7D, a one-way ANOVA was performed, with ** = p value < 0.01. For Figure 7E, significance was determined using a Mantel-Cox test with Bonferroni correction; * = p value 0.05. One-way ANOVA was used determine statistical significance between treatment groups in Figures 7F, with * = p value < 0.05, ** = p value < 0.01, and *** = p value <0.001. Figure 7D-E representative of n=2 similar studies. See also Figure S11.

During evaluation of meso eCAR-T activity in vivo in pilot studies, we observed that dual-edit meso eCAR-Ts drove body weight loss likely due to reactivity of endogenous TCRs for xenoantigens. Thus, we included a SOCS1/Regnase-1 /TRAC triple-edited meso eCAR-T group (Figure S11D-E). Upon transfer of 7×10^5^ meso eCAR-Ts, tumor growth inhibition was observed by both SOCS1 and Regnase-1 single edit meso eCAR-Ts, and with dual and triple edited cells driving tumor regression (Figure 7D, individual mice in Figure S11F). Triple edited meso eCAR-Ts were able to significantly enhance survival in comparison to both single and dual edits as well as controls (Figure 7E). Examination of meso eCAR-T frequency in the peripheral blood demonstrated accumulation of Regnase-1 and SOCS1 single, dual and triple edited meso eCAR-Ts (Figure 7F), with dual-edited eCAR-T showing the greatest level of accumulation at Day 28 and corresponding with body weight loss, suggestive of xeno GvHD driven by the dual-edited meso eCAR-Ts (Figure S11G). No body weight loss was observed with triple edited meso eCAR-Ts. Upon study end, strong tumor infiltration of both dual and triple edited meso eCAR-Ts was observed (Figure 7G). Lastly, to examine the contribution of exogenous cytokines to meso eCAR-T efficacy, meso eCAR-Ts were evaluated in NSG and NSG-Tg(Hu-IL-15) mice. Control meso eCAR-Ts displayed a modest increase in anti-tumor activity in the NSG-Tg(Hu-IL15) strain, presumably driven by increased availability of IL-15. In contrast, dual-edited meso eCAR-Ts displayed similarly strong anti-tumor activity in both NSG and NSG-Tg(Hu-IL15) strains (Figure S11H). This indicates strong anti-tumor activity of dual-edited meso eCAR-Ts independent of exogenous IL-15 support. These data collectively demonstrate that inactivation of SOCS1 and Regnase-1 in human T cell therapies cooperate to enhance anti-tumor functionality against solid tumors.

## DISCUSSION

In this study, we provide a genome-wide functional roadmap of the key genes and pathways regulating CD8^+^ T cell function against solid tumors. Negative regulators of the NF-κB and cytokine signaling pathways were identified as the strongest brakes on CD8^+^ T cell function, with multiple novel targets additionally identified and warranting further exploration. Regnase-1 and SOCS1 were identified as top hits in the genome, with Regnase-1 implicated as impacting CD8^+^ T cell function through the NF-κB pathway, and SOCS1 a known negative regulator of cytokine signaling. Our combination screens reveal the criticality of NF-κB signaling in CD8^+^ T cell function by identifying neutralizing interactions between Regnase-1, Roquin-1, Traf6, IκΒα, A20 and Peli1. Neutralizing interactions between Traf6 with Roquin-1, Regnase-1, IκΒα, Peli1 and A20 demonstrate pathway redundancy and implicate Traf6 as upstream of these regulators. Regnase-1 and Roquin-1 did not exhibit neutralizing interactions indicating complementarity rather than redundancy, which is consistent with a prior report showing dual-edit driven enhancement of CAR-T activity in pre-clinical models[59,62]. Our combination results additionally pinpointed negative regulators of NF-κB and cytokine signaling as promising dual-edit pairs, and evaluation of multiple combinations identify SOCS1/Regnase-1 dual-edits as driving the strongest anti-tumor functionality. In addition to optimizing functional enhancement, an added benefit of combining negative regulators from two different pathways is the potential to avoid unconstrained biology driven by pathway hyperactivation, such as the toxicities seen with dual-inactivation of Regnase-1 together with Roquin-1 in pre-clinical CAR-T models due to uncontrolled outgrowth[59].

Wei et al have previously reported Regnase-1 as a regulator of CD8^+^ T cell anti-tumor function, with SOCS1 implicated as a combination partner[26,28]. In addition to our genome-wide and combination screen insights detailed above, our study confirms and extends this work in multiple ways. First, we provide evidence that Regnase-1 inactivation may be the most effective edit capable of enhancing CD8^+^ T cell anti-tumor function, and with dual-inactivation of SOCS1 and Regnase-1 driving the strongest enhancement amongst all evaluated dual-edits. Importantly, we detail how SOCS1 and Regnase-1 inactivation impacts Tex and memory subset differentiation and function and extend these insights to develop novel human T cell therapeutics.

We define how SOCS1 and Regnase-1 inactivation perturbs the trajectory and function of tumor-specific CD8^+^ T cells along the Tex and memory differentiation continuum. Regnase-1 inactivation broadly enhanced the accumulation of Tex subsets, and facilitated the formation of persistent T_em_ cells following tumor clearance. SOCS1 inactivation conversely enhanced the accumulation of Tex^int^ and Tex^eff^ cells within tumor while driving the formation of persistent T_cm_ cells, consistent with our prior work[31] as well as studies reporting IL-2 can drive similar effector cell increases[63–65]. The strong enhancement in anti-tumor activity observed by SOCS1 and Regnase-1 dual inactivation is mediated through the combined effects of each single edit strongly expanding Tex^int^ cells in blood and spleen, Tex^int^ and Tex^eff^ cells in TDLN, and Tex^eff^ cells in tumor. Dual inactivation rewired the transcriptional status of Tex and memory subsets by increasing the expression of effector regulators and Regnase-1 substrates while decreasing the expression of transcriptional regulators of terminal exhaustion including TOX, as well as BIM. BIM expression has recently been reported to be suppressed by Batf[66], suggestive of a Regnase-1/Batf/BIM regulatory circuit warranting further investigation. The observed fewer Tex^prog1^ cells in the dual-edit versus Regnase-1 single edit may be a result of increased SOCS1 inactivation-driven cytokine sensitivity by this subset, lowering the threshold for differentiation. Lastly, and consistent with our combination-informed approach in targeting non-overlapping pathways, we observed little overlap between SOCS1 and Regnase-1 -driven transcriptional signatures, with the dual-edited signature a composite in both OT1s and eTIL. We propose that when considering combination approaches for therapeutic intervention, targeting non-overlapping and mechanistically complementary pathways should be considered to maximize functional impact. To conclude, dual-inactivation of SOCS1 and Regnase-1 combine to strongly enhance the accumulation of Tex and memory cell subsets optimized for effector function.

Our analysis also evaluated the impact of Regnase-1 inactivation as a single edit as well as in combination with SOCS1 on memory cell differentiation. Regnase-1 inactivation drove the accumulation of a population of Tem^Gzmk^ and T_memory_ cells within tumor, consistent with Wei et al reporting Regnase-1 inactivation facilitating effector memory formation[28]. Following tumor clearance, Regnase-1 inactivation facilitated the formation of CD8^+^ T cells displaying a T_em_ state which persisted at low overall frequencies in blood versus other edits. Combining SOCS1 inactivation with Regnase-1 maintained a similar T_em_ state while increasing the overall frequency of persistent cells. The ability of SOCS1 to ‘rescue’ the Regnase-1-driven frequency deficit following tumor clearance highlights the benefits of targeting non-overlapping and complementary pathways. Indeed, a dual-edit of Regnase-1/BCOR has been described as facilitating an immortal stem-like state in CD8^+^ T cells [67,68]. We note commonalities between our findings and Wang et al., including the frequency deficit of persisting Regnase-1 single-edited memory CD8^+^ T cells, which can be rescued by the addition of a second edit. There are, however, clear differences in the transcriptional programs driven between SOCS1 and BCOR. SOCS1 drives increased STAT signaling facilitating increased overall survival, differentiation into effector cells at the sites of antigen engagement and increased effector function [31], whereas BCOR inactivation locks Regnase-1 inactivated CD8^+^ T cells into a core stemness transcriptional program [67].

Our data exploring the therapeutic potential of SOCS1/Regnase-1 dual-edits in eTIL and eCAR-Ts demonstrate translation of our preclinical mouse observations to human therapeutic candidates. In mouse systems encompassing mTIL and OT1s, Regnase-1 exhibited a more pronounced impact compared to SOCS1. In human-derived eTIL, the opposite was observed, with SOCS1 having a greater impact in comparison to Regnase-1. It is likely that these differences are driven by different experimental settings: human TIL are known to be primarily derived from memory cells with a significant cellular age and following ex vivo expansion over weeks, while murine OT1s and mTIL are derived from a younger cellular provenance and expanded ex vivo over days. Importantly, we observed imposition of both SOCS1 and Regnase-1-driven signatures across both mouse and human systems, and in eTIL, KSQ-004EX possessed a composite of SOCS1 and Regnase-1 single-edit signatures. Limitations of this study includes the need for additional investigation as to how Regnase-1 may impacts NF-κB signaling in Tex cells, with induction of the transcriptional regulator *Nfkbiz*, encoding for IκBζ, a candidate mechanism. Indeed, increased IκBζ- driven transcriptional activity was identified as being mediated by Regnase-1 inactivation in our eTIL transcription factor analyses. These data together with the refractoriness of KSQ-004EX to chronic stimulation and the strong in vivo combination effect by dual-edited meso eCAR-Ts support the evaluation of SOCS1 and Regnase-1 dual-inactivation in human T cell therapies. Collectively, this study describes the mechanisms as to how inactivation of SOCS1 and Regnase-1 maximizes CD8^+^ T cell anti-tumor functionality for use in patients with refractory solid tumors. KSQ-001EX and KSQ-004EX are actively being evaluated in a Phase 1/2 clinical trial (NCT06237881 and NCT06598371) for the treatment of refractory and metastatic solid tumors.

## Supporting information

Graphic Abstract

## SUPPLEMENTAL INFORMATION

**Document S1. Figures S1–S11**

**Document S2. Table S1**

**Document S3. Table S2**

**Document S4. Table S3**

**Document S5. Table S4**

**Document S6. Table S5-S7**

**Document S7. Table S8-S10**

**Document S8. Table S11**

## DECLARATIONS

### Ethics approval and consent to participate

This study was conducted in accordance with the Declaration of Helsinki ethical guidelines. This study was approved by the Dana Farber Institutional Review Board (DF/HCC# 09-472) and informed consent was obtained from participants who provided tumor tissue for research.

### Availability of data and material

All unique/stable reagents generated in this study are available from the lead contact upon reasonable request for non-commercial use with a completed materials transfer agreement. RNA-Seq data from Figures 3 and 6 and scRNA-Seq data from Figures 4 and 5 are available via NCBI Gene Expression Omnibus (GEO) database and are publicly available as of the date of publication at GSE299241 and GSE299242. Data reported in this paper will be shared by the lead contact upon reasonable request. This paper does not report original code. Any additional information required to reanalyze the data reported in this paper is available from the lead contact upon request.

### Competing Interests

All KSQ Therapeutics and CTMC affiliated authors are current or former employees as well as shareholders of their respective affiliations. Chantale Bernatchez is a member of KSQ Therapeutics Scientific Advisory Board. Frank Stegmeier is a member of the KSQ Therapeutics Board of Directors. Glenn Hanna received Sponsored Research Agreement (SRA) funding from KSQ Therapeutics in support of this work. Patent applications and granted patents from this work include WO2019/178420, WO2019/178421, WO2020/163365, WO2021/108455, and WO2025106513.

### Funding

All work was funded by KSQ Therapeutics. KSQ therapeutics didn’t influence the results/outcomes of the study despite author affiliations with the funder.

### Authors’ Contributions

Conceptualization: I.L.M, D.M, K. H-V, K.W, G.M, G.H., G.K., L.C., M-A. F., C.B., J.G., F.M, D.S., M.S. and M.J.B. Investigation: I.L.M., D.M., K. H-V., A.D., D.M., B.H., A.Y., S.M., S.L., N.P., A.S., P.D, M.G., C.C., H.G., S.J., F.T., S.C., E.F., G.K, J.M., C.B., C.W., L.C., and M.S. Supervision: K.W., F.S., J.G., F.M., D.S., M.S and M.J.B. Writing: M.J.B. Equal co-first authorship contributions determined as I.L.M conceived of and performed CRISPR screens depicted in Figure 1-2 and mTIL model in Figure 3; D.M. and K.H-V. performed experiments depicted in Figure 4-5, and K.W oversaw the development and characterization of KSQ-004EX eTIL.

## Acknowledgements

Figures 1A, 2A, S3A and the graphical abstract were created with BioRender. We’d like to thank Dr. Jonathan Weissman for generously reviewing and commenting on the manuscript.

## Authors’ Information

Requests for further information and resources should be directed to and will be fulfilled by the lead contact, Dr. Micah J. Benson.

## MATERIALS AND METHODS

### REAGENT & RESOURCE TABLE

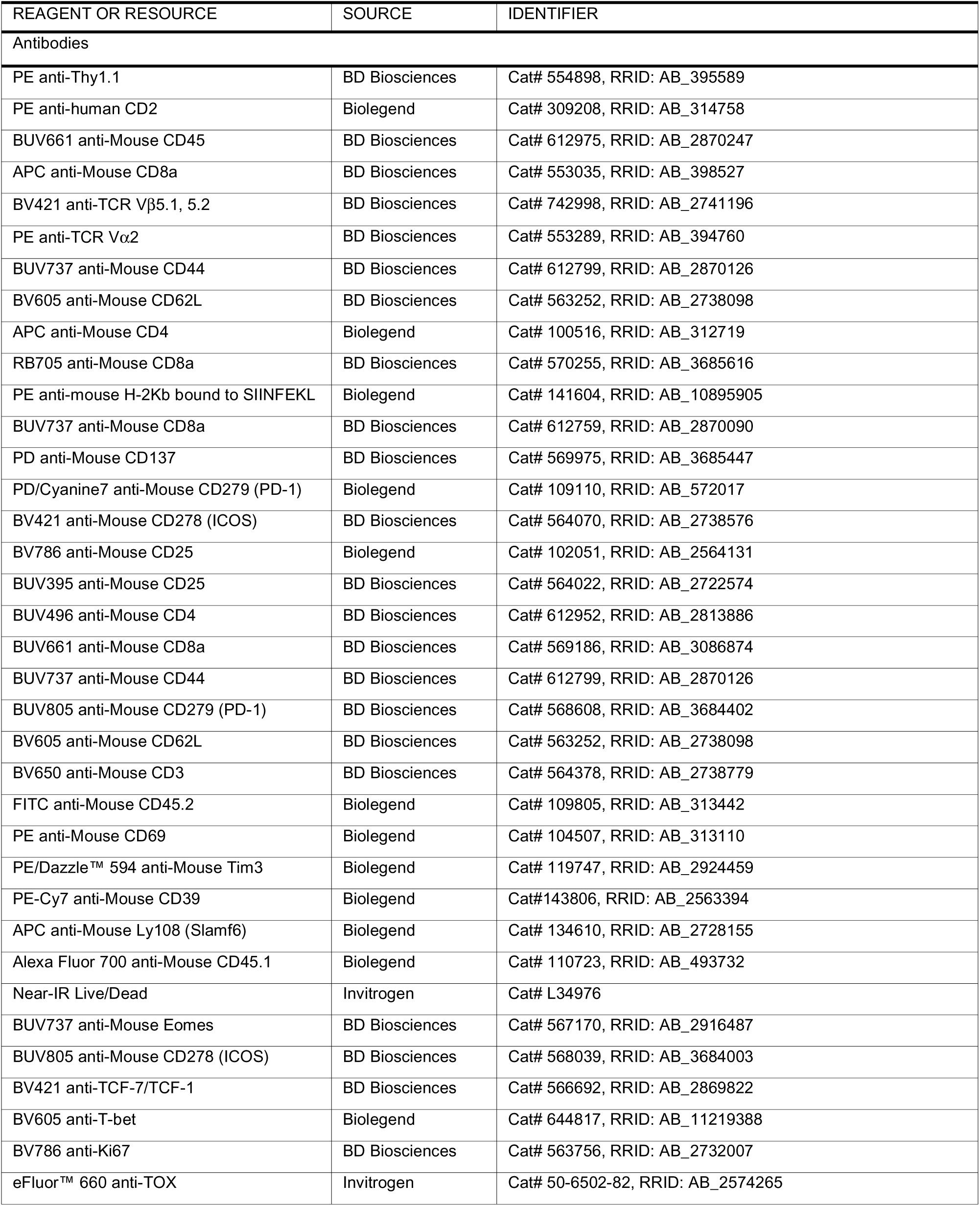
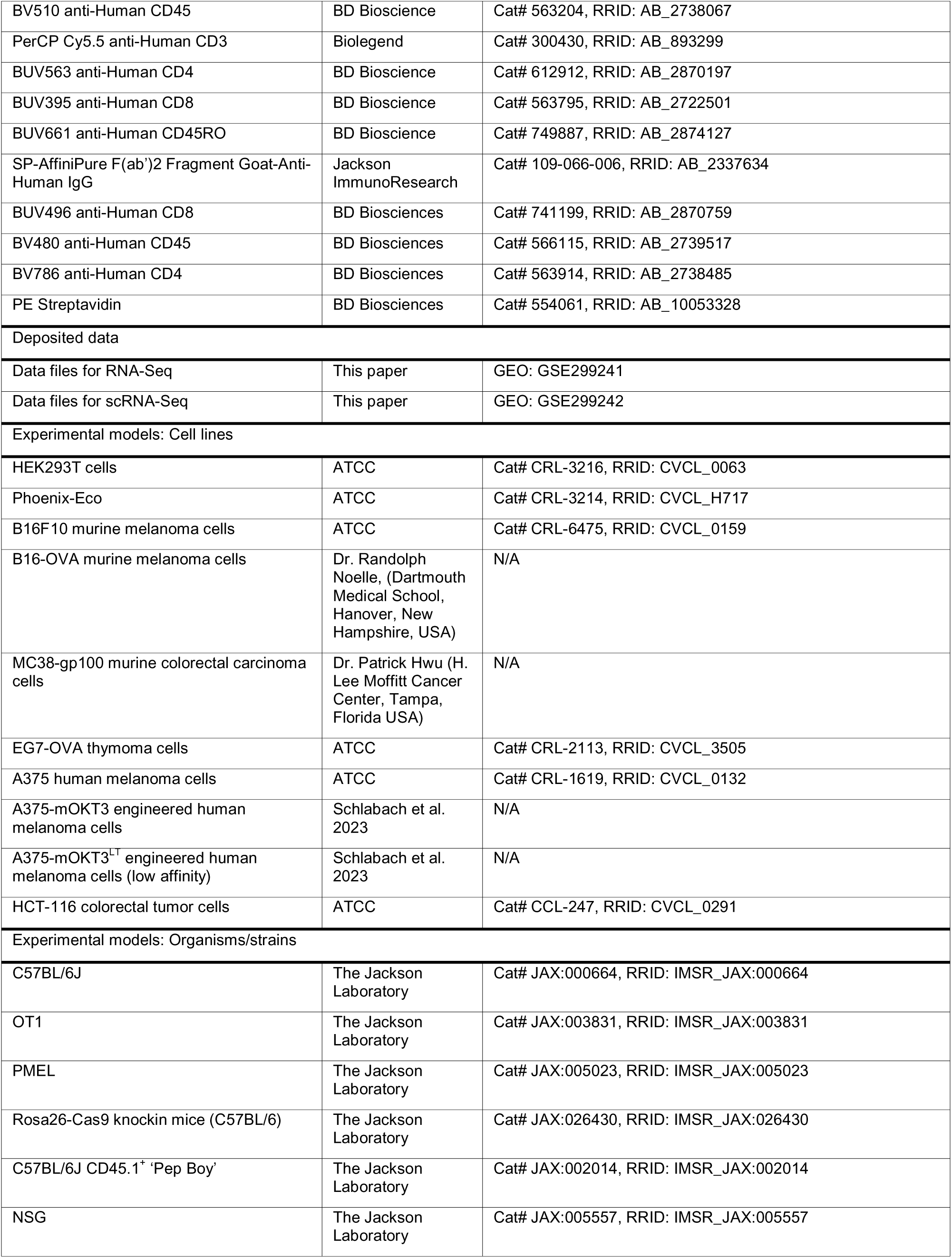
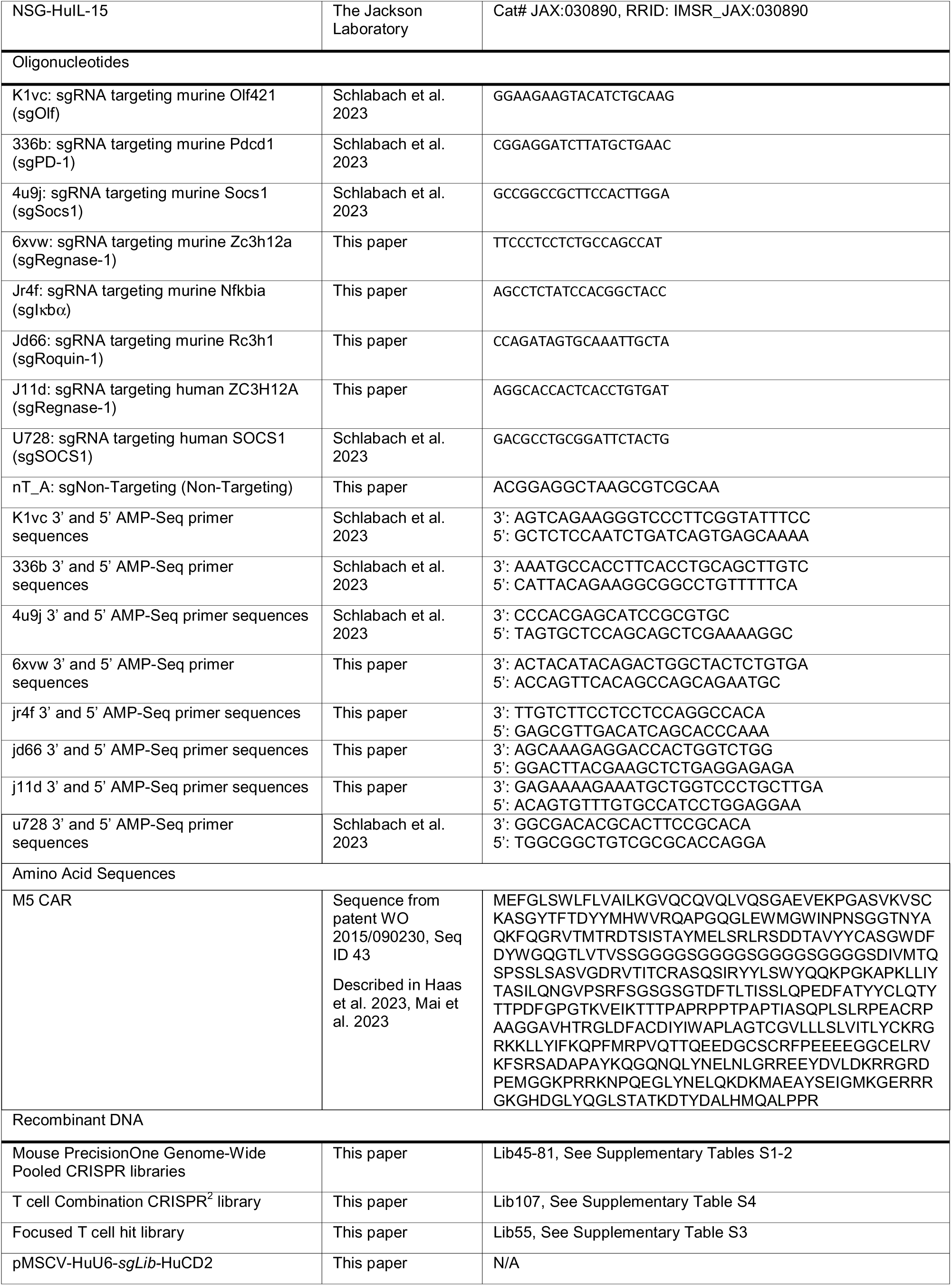
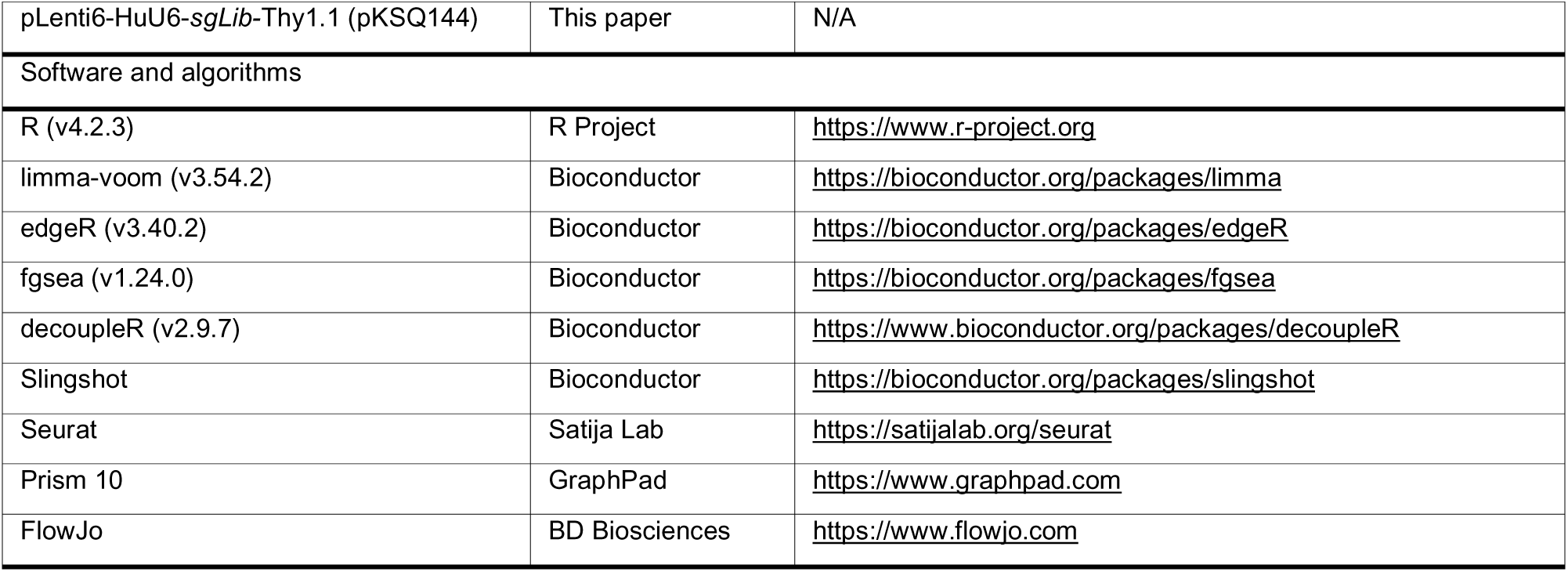

## EXPERIMENTAL MODEL DETAILS

### Mice

Six-to-nine week old female C57BL/6J, OT1, PMEL, NSG, NSG-HuIL-15, and CD45.1^+^ mice were purchased from The Jackson Laboratory. To generate Cas9-Tg x TCR-Tg strains for in vivo CRISPR screens, female Cas9 mice were crossed with OT1 or PMEL mice at The Jackson Laboratory. Mice were injected with tumors between 6-12 weeks of age. All procedures involving the care and use of animals were reviewed and approved by the Institutional Animal Care and Use Committee (IACUC) and were conducted in the KSQ vivarium under protocol NC-053 in accordance with associated regulations and guidelines.

### Cell Lines

The B16-OVA cell line was kindly provided by Dr. Randolph Noelle (Dartmouth Medical School, Hanover, NH), and the MC38-gp100 colon cell line kindly provided by Dr. Patrick Hwu (H. Lee Moffitt Cancer Center, Tampa, FL). B16F10, EG7-OVA and HCT-116 were purchased from ATCC. The A375 human melanoma cell line was obtained from ATCC and engineered to express either a low-affinity or high-affinity membrane associated anti-CD3 binding domain from clone OKT3 (mOKT3) together with RFP as previously described (Schlabach et al. 2023), with the low-affinity A375-mOKT line used for in vitro spheroid cytotoxicity and IFN- release assays, and the high-affinity A375-mOKT3 line used for the chronic stimulation assay[31]. B16-Ova, MC38-gp100, HCT-116 and A375-mOKT3 cells were cultured in DMEM (Gibco, Cat# 11885076) supplemented with 10% FBS (Gibco, Cat# 10082147). EG7-OVA cells were cultured in RPMI 1640 medium supplemented with 2 mM L-glutamine, 1.5g/l sodium bicarbonate, 4.5 g/l glucose, 10 mM HEPES, 1 mM sodium pyruvate, 0.05 mM 2-BME, 0.4 mg/ml G418, 10 % FBS. Cells were passaged two to three times per week to ensure confluency in the flask never surpassed 80%. Tumor cells were harvested in serum-free DMEM during exponential growth phase immediately prior to inoculation.

### Engineering of mouse primary TCR-Tg CD8^+^ T cells

Spleens were harvested from OT1 or PMEL mice, dissociated using a GentleMACS octo dissociator and CD8 T cells isolated using the EasySep^TM^ Mouse CD8+ T Cell Isolation Kit (Stemcell, Cat# 19853) according to the manufacturer’s instructions. Purified CD8^+^ T cells were activated with mouse CD3/CD28 Dynabeads and 4 ng/ml mouse rIL-2. 48 hours later, beads were removed and sgRNA / Cas9 RNP complexes prepared with sgRNA at 22 mM and Cas9 at 15 mM in IDTE, Buffer T and electroporation enhancer. CD8 T cells were electroporated at 1700V, 20ms, 1pulse using a Neon transfection system (LifeTechnologies). Cells were further expanded with 32 ng/ml mouse rIL-2 for 48 hours, harvested, and either transferred directly into recipient mice or cryopreserved in 90% FBS + 10% DMSO. Engineered cells were evaluated for editing efficiency of the target gene and tested via PCR for a comprehensive list of mouse pathogens by Charles River Laboratory Testing Management prior to infusion.

### OT1 / B16-Ova Tumor Model

Female C57BL/6J mice were inoculated with 0.5×10^6^ B16-Ova cells subcutaneously in the right flank. After eight days for small tumor studies (∼100mm^3^; range of 80mm^3^-120mm^3^) or 14 days for large tumor studies (mean 350mm^3^; range of 135mm^3^-550mm^3^) mice were randomized by tumor volume and 3×10^6^ edited OT-1 T cells were transferred in 200µl PBS via the lateral tail vein. Tumor volumes were measured using a digital caliper and tumor volume (mm3) was calculated using the following formula (width^2^ x length) *0.5, where length is the longer dimension. To assess tumor growth following re-challenge in Figure S4 and S8, mice undergoing complete tumor rejection were re-challenged with 0.5×10^6^ B16-Ova cells subcutaneously on the left flank on the indicated day and monitored for tumor growth.

### mTIL / B16-Ova Tumor Model

Murine CD3^+^ TIL (CD4/CD8) were isolated from B16-OVA subcutaneous tumors from 120 donor mice on a CD45.1 background using CD4/CD8 MicroBeads (Miltenyi cat#130-116-480). After isolation, mTIL were cultured in complete T cell media with 3,000 IU/mL human IL-2 for 3 days prior to electroporation (EP) using the Neon transfection system (LifeTechnologies) with CRISPR associated protein 9 (Cas9) ribonucleoprotein (RNP) complexes. Edited mTIL were then cultured in the presence of 3,000 IU/mL IL-2 for an additional 5 days prior to characterization and adoptive cell transfer. Target editing efficiencies for mTIL in Figure 3F were sgOlf: 67%; sgSocs1: 87%; sgRegnase-1: 86%; and sgSocs1 + sgRegnase-1: 82% and 79%. On Day 0, CD45.2 recipient mice bearing B16-OVA subcutaneous tumors (95 mm^3^) were dosed with 7×10^6^ edited mTIL (n = 10 mice/group). Tumor volumes were measured twice weekly.

### PMEL / Disseminated B16F10 Tumor Model

Mice were injected i.v. with 0.5×10^6^ B16F10 cells, with 10×10^6^ edited PMEL T cells injected i.v. on day 3 following tumor injection. For Figure 3c, two or three mice per group were examined for lung metastasis burden on Day 15 as indicated. Survival was used as the primary endpoint in Figure 3d, with n=5 mice per treatment group.

### Meso eCAR-T / HCT-116 Tumor Model

Generation of Mesothelin CAR-Ts: PBMC were isolated from leukopaks (Discovery Life Sciences) using the EasySep Direct Human PBMC Isolation Kit (StemCell, Cat# 19654) and cryopreserved. Upon thaw into HABS (Valley Biomed, Cat# HP1022Hi) T cells were purified using the EasySep Human T Cell Isolation Kit (StemCell, Cat# 17951) and activated for 3 days using T Cell TransAct (Miltenyi Biotec, Cat# 130-111-160) in X-VIVO 15 (Lonza, Cat# BP04-744Q) with 5% HABS, 10 µM of HEPES (Gibco, Cat# 15630080), 1x GlutaMAX (Gibco, Cat# 35050-31), and 1,000 IU/mL of IL-2 (Miltenyi Biotec, Cat# 130-097-748) (‘T cell base media’). On day 1, T cells were transduced with lentivirus containing the M5 CAR tagged with RQR8 by spinfection using non-TC treated 6 well plates coated with retronectin (Takara, Cat# T100B). Cells were incubated overnight, virus was removed, and CAR-Ts were electroporated using the same protocol as described for TIL. For triple KO CAR-Ts, 1.8 µM of each gRNA was used. Knockout efficiency was determined by AMP-Seq and transduction efficiency assessed by flow cytometry using a CD34 antibody (ThermoFisher, Cat# MA1-10205). CAR-Ts were expanded for a total of 8-10 days and cryopreserved using CryoStor CS10 (StemCell, Cat# 07930). A total of 5 × 10L HCT-116 cells, suspended in 200 µl of saline, were injected subcutaneously into the right flank of NSG or NSG-IL15 transgenic mice bearing tumors with an average volume of approximately 100 mm³. Tumor-bearing mice were randomized into treatment groups (n = 5–6 per group). On Day 0 of the study, each mouse received an i.v. injection of 0.7 × 10L CAR+ T cells. A separate control group (n = 5) received untransduced T cells equivalent to the total T cells administered in the CAR+ T cell groups. Tumor volumes and body weights were monitored twice weekly using calipers and scales, respectively. For ex vivo enumeration of Meso eCAR-T Cells, blood was collected via submandibular survival bleeds from mice in each treatment group post-CAR-T infusion. Following RBC lysis, cell pellets were incubated with mouse Fc receptor (FcR) block (BioLegend, catalog no. 101320), Human TruStain FcX (BioLegend) and biotinylated SP-AffiniPure F(ab’)² Fragment Goat Anti-Human IgG (Jackson ImmunoResearch). Quantification and immunophenotyping of CAR-T cells were performed using a flow cytometric panel with antibodies as detailed. CAR^+^ T cells were defined as the CD45^+^CAR^+^ population. Quantification was conducted using CountBright™ absolute counting beads (Thermo Fisher Scientific); On Day 42, remaining study mice were euthanized, and spleens and tumors were harvested for FACS analysis.

## METHOD DETAILS

### CRISPR Screens

Libraries were designed as followed:

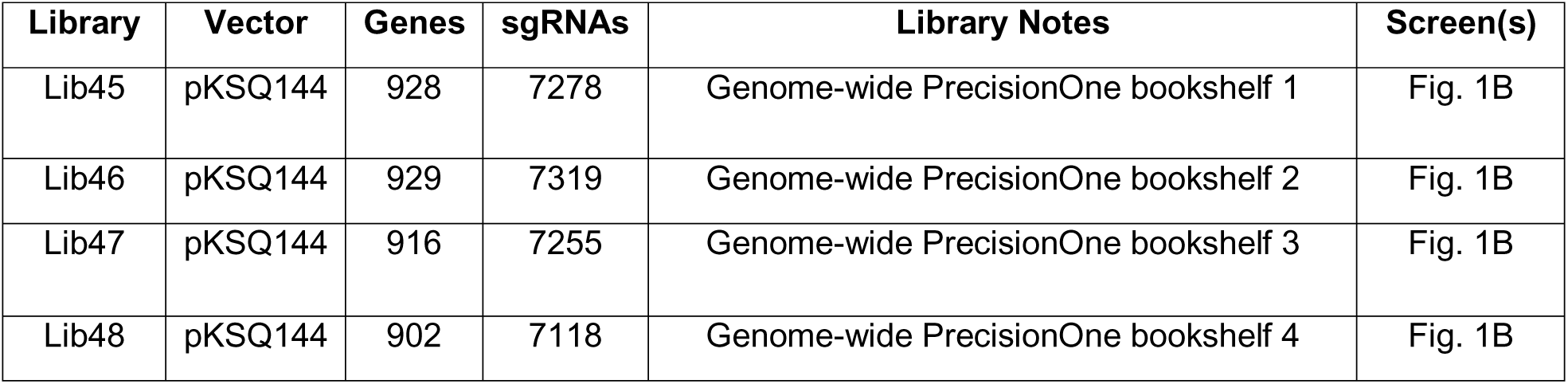
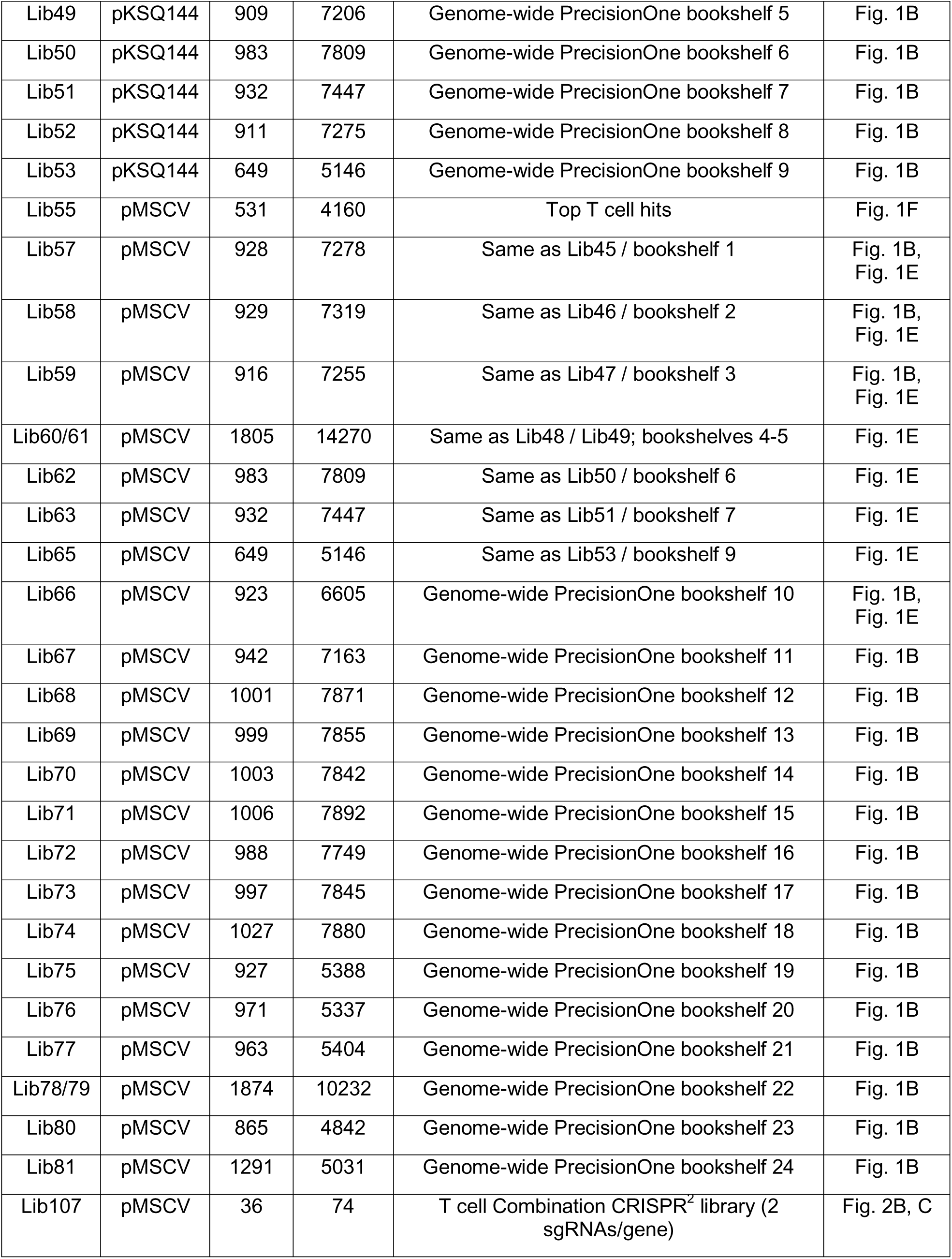

| Library | Vector | Genes | sgRNAs | Library Notes | Screen(s) |
| --- | --- | --- | --- | --- | --- |
| Lib45 | pKSQ144 | 928 | 7278 | Genome-wide PrecisionOne bookshelf 1 | Fig. 1B |
| Lib46 | pKSQ144 | 929 | 7319 | Genome-wide PrecisionOne bookshelf 2 | Fig. 1B |
| Lib47 | pKSQ144 | 916 | 7255 | Genome-wide PrecisionOne bookshelf 3 | Fig. 1B |
| Lib48 | pKSQ144 | 902 | 7118 | Genome-wide PrecisionOne bookshelf 4 | Fig. 1B |
| Lib49 | pKSQ144 | 909 | 7206 | Genome-wide PrecisionOne bookshelf 5 | Fig. 1B |
| Lib50 | pKSQ144 | 983 | 7809 | Genome-wide PrecisionOne bookshelf 6 | Fig. 1B |
| Lib51 | pKSQ144 | 932 | 7447 | Genome-wide PrecisionOne bookshelf 7 | Fig. 1B |
| Lib52 | pKSQ144 | 911 | 7275 | Genome-wide PrecisionOne bookshelf 8 | Fig. 1B |
| Lib53 | pKSQ144 | 649 | 5146 | Genome-wide PrecisionOne bookshelf 9 | Fig. 1B |
| Lib55 | pMSCV | 531 | 4160 | Top T cell hits | Fig. 1F |
| Lib57 | pMSCV | 928 | 7278 | Same as Lib45 / bookshelf 1 | Fig. 1B,<br>Fig. 1E |
| Lib58 | pMSCV | 929 | 7319 | Same as Lib46 / bookshelf 2 | Fig. 1B,<br>Fig. 1E |
| Lib59 | pMSCV | 916 | 7255 | Same as Lib47 / bookshelf 3 | Fig. 1B,<br>Fig. 1E |
| Lib60/61 | pMSCV | 1805 | 14270 | Same as Lib48 / Lib49; bookshelves 4-5 | Fig. 1E |
| Lib62 | pMSCV | 983 | 7809 | Same as Lib50 / bookshelf 6 | Fig. 1E |
| Lib63 | pMSCV | 932 | 7447 | Same as Lib51 / bookshelf 7 | Fig. 1E |
| Lib65 | pMSCV | 649 | 5146 | Same as Lib53 / bookshelf 9 | Fig. 1E |
| Lib66 | pMSCV | 923 | 6605 | Genome-wide PrecisionOne bookshelf 10 | Fig. 1B,<br>Fig. 1E |
| Lib67 | pMSCV | 942 | 7163 | Genome-wide PrecisionOne bookshelf 11 | Fig. 1B |
| Lib68 | pMSCV | 1001 | 7871 | Genome-wide PrecisionOne bookshelf 12 | Fig. 1B |
| Lib69 | pMSCV | 999 | 7855 | Genome-wide PrecisionOne bookshelf 13 | Fig. 1B |
| Lib70 | pMSCV | 1003 | 7842 | Genome-wide PrecisionOne bookshelf 14 | Fig. 1B |
| Lib71 | pMSCV | 1006 | 7892 | Genome-wide PrecisionOne bookshelf 15 | Fig. 1B |
| Lib72 | pMSCV | 988 | 7749 | Genome-wide PrecisionOne bookshelf 16 | Fig. 1B |
| Lib73 | pMSCV | 997 | 7845 | Genome-wide PrecisionOne bookshelf 17 | Fig. 1B |
| Lib74 | pMSCV | 1027 | 7880 | Genome-wide PrecisionOne bookshelf 18 | Fig. 1B |
| Lib75 | pMSCV | 927 | 5388 | Genome-wide PrecisionOne bookshelf 19 | Fig. 1B |
| Lib76 | pMSCV | 971 | 5337 | Genome-wide PrecisionOne bookshelf 20 | Fig. 1B |
| Lib77 | pMSCV | 963 | 5404 | Genome-wide PrecisionOne bookshelf 21 | Fig. 1B |
| Lib78/79 | pMSCV | 1874 | 10232 | Genome-wide PrecisionOne bookshelf 22 | Fig. 1B |
| Lib80 | pMSCV | 865 | 4842 | Genome-wide PrecisionOne bookshelf 23 | Fig. 1B |
| Lib81 | pMSCV | 1291 | 5031 | Genome-wide PrecisionOne bookshelf 24 | Fig. 1B |
| Lib107 | pMSCV | 36 | 74 | T cell Combination CRISPR<sup>2</sup> library (2 sgRNAs/gene) | Fig. 2B, C |

To construct libraries for mouse T cell screening, libraries were cloned into either a pLenti6-based lentiviral backbone containing a human U6 promoter cassette with a Thy1.1 selectable marker (pKSQ144) or a self-inactivating murine stem cell virus (pMSCV) retroviral vector backbone also containing a human U6 promoter cassette with a human CD2 marker. Vectors additionally contained a library of randomly synthesized 12-mer barcodes, or unique molecular identifiers (UMIs), with the nucleotide sequence BVHDBVHDBVHD (IUPAC mixed base codes). These UMIs (theoretical diversity of 531,441 unique sequences), when randomly paired with the guide sequence during cloning, allow the ability to independently trace millions of unique infection events in immune cells. Libraries were generated by searching for PAM sites (NGG) in the human and mouse genome that were expected to correspond to sgRNAs that would cut the coding sequence of genes of interest. sgRNAs were chosen for the library that were predicted to cut the genome in at most one site, while not overlapping with known high prevalence SNPs and having few (<10) closely related sites in the genome that could represent likely off-target sites. Several classes of controls were included, such as non-cutting, olfactory receptor cutting not expected to impact biology in immune cells, and lethal shredders that cut the genome thousands of times. The final set of filtered sgRNAs were then synthesized on a microarray (Agilent oligo library synthesis) as oligonucleotides with adapters, PCR amplified and cloned using the type IIs restriction enzyme BbsI into the final lentiviral or retroviral vector.

### sgRNA Library Virus Production

Lentivirus was generated by lipid transfection of packaging plasmids into HEK293T cells. For a 10-layer CellStack (Corning), 300 million HEK239T cells were plated in 1 L of DMEM + 10% FBS. 24 hours later transfection was performed using 461 ug of sgRNA library plasmid, 231 µg of packaging plasmid psPax2 (Gag-Pol), and 115 µg of pMD2.G packaging plasmid (VSV-G). Plasmid DNA was added to 37.77 ml of room temperature OptiMEM media. 2428 µl of TransIT transfection reagent (Mirus Bio) was added to this mixture, mixed by vortexing, and incubated for 20 min before proceeding. Transfection mixture was added to 1 L of DMEM + 10% FBS. Media was aspirated from HEK293T cells and fresh media containing transfection mixture was added. 24 hours after transfection the media was replaced 1 L of UltraCulture media (Lonza) supplemented with L-glutamine. 48 hours after transfection the supernatant was harvested from the cells, incubated for 1 hour at 37°C in the presence of 50 units/ml benzonase, filtered using a 0.45 µM bottle top PES filter, and concentrated by tangential flow filtration on a spectrum labs KrosFlow mPES hollow fiber filter (100kD molecular weight cutoff). Virus was concentrated approximately 10-50x, depending on initial volumes being processed.

Retrovirus was generated by transient transfection of Phoenix-Eco retroviral packaging cells with library plasmid pools. 5×10^6^ Phoenix-Eco cells were plated per well 24 hours before transfection in DMEM + 10% FBS. Transfection is carried out by mixing 10 µg of sgRNA library plasmid with 327 µl optiMEM media. 21 µl of Mirus TransIT-293 transfection reagent is added to this mixture, vortexed briefly, and incubated for 20 min. Transfection mixture is then added dropwise to previously plated phoenix cells and incubated, with media replaced after 24 hours with 5 ml of RPMIc (RPMI + 10% HI FBS, 20 mM HEPES, 50 µM 2-mercaptoethanol). Virus was frozen and titered for subsequent large-scale transduction.

### Engineering of mouse TCR-Tg CD8^+^ T cells

Cas9-Tg x TCR-Tg OT1 or PMEL CD8 T cells were isolated from freshly harvested mouse spleens and dissociated using a GentleMACS system (Miltenyi), and with CD8 T cells purified by negative selection (EasySep Mouse CD8+ T cell isolation kit). CD8s were placed in T225 flasks at a concentration of 1×10^6^ cells/ml and activated with CD3/CD28 Dynabeads and 2 ng/ml mouse rIL-2 with cRPMI. The following day, T cells were transduced wither either lentivirus or retrovirus. Dynabeads were removed on Day 3, with cells resuspended on cRPMI + 32ng/ml IL-2. On Day 5, transduced T cells were harvested and selected by positive selection using either Thy1.1 microbeads for lentivirus or hCD2 microbeads for retrovirus and cryopreserved for future use.

### OT1 / B16-OVA genome wide screen

Following subcutaneous tumor inoculation in the right flank with 0.5×10^6^ cells, mice were randomized on Day 12 with a median tumor size ranging between 108mm^3^-120mm^3^, depending on bookshelf. 5×10^6^ Thy1.1^+^ or hCD2^+^ Cas9-Tg x OT1 CD8 T cells per mouse were injected i.v., with 7-8 mice per bookshelf. Mice were euthanized and tissues harvested 14 post cell transfer.

### PMEL / MC38-gp100 screen

Following subcutaneous tumor inoculation in the right flank, mice were randomized on Day 7 with a median tumor size of 85mm^3^. 7×10^6^ hCD2^+^ Cas9-Tg x PMEL CD8 T cells were injected i.v., with 8 mice per bookshelf. Mice were euthanized and tissues harvested 14 days post cell transfer.

### OT1 / B16-OVA Lib55 screen

Following subcutaneous B16-Ova tumor inoculation in the right flank, mice were randomized on Day 12, with 6 mice receiving 5×10^6^ hCD2^+^ Cas9-Tg x OT1s. Mice were euthanized and tissues harvested 14 days post transfer.

### OT1 / EG7-OVA Lib55 screen

Following subcutaneous inoculation of mice with 1×10^6^ cells on the right flank, mice were randomized on Day 8 with median tumor size of 168mm^3^. 8 mice received 5×10^6^ hCD2^+^ Cas9-Tg x OT1s. Mice were euthanized, and tissues harvested 11 days post transfer.

### OT1 / B16-OVA Combination screen

Mice were randomized on Day 7 following tumor inoculation, with median tumor size of 193mm^3^, with 3×10^6^ hCD2^+^ Cas9-Tg x OT1 cells transferred. Mice were euthanized and tissues harvested 14 days following initial OT1 transfer.

### PMEL / MC38-gp100 Combination screen

Mice were randomized on Day 7 following tumor inoculation, with median tumor size of 92mm^3^, with 7×10^6^ hCD2^+^ Cas9-Tg x PMEL transferred. Mice were euthanized and tissues harvested 14 days following initial PMEL transfer.

For all screens, 10×10^6^ CD8^+^ T cells were saved to determine the sgRNA distribution of the input population of T cells. At the indicated time following adoptive transfer of cells, mice were euthanized and blood was harvested in EDTA tubes, and stored at −80°C. Tumors, spleens, tumor draining and non-draining lymph nodes were harvested and processed further for enrichment of CD8 T cells using CD8a Microbeads (Miltenyi, cat# 130-049-401) from the spleen and CD45 Microbeads (Miltenyi, cat# 130-052-301) from the tumor. Tumors were digested using the Miltenyi Tumor Dissociation Kit (Cat# 130-096-730) according to the manufacturer’s instructions. Genomic DNA was isolated using the Qiamp Blood Midi and Maxi kids, according to the manufacturer’s instructions.

### CRISPR Screen Analysis

Paired-end sequencing of the sgRNA position(s) and UMI position was performed to determine sgRNA and clone identity, respectively. Counts of sgRNA frequency were generated from FASTQ sequencing files by counting the number of occurrences of each sequence at the expected sgRNA position. For each sgRNA sequence, counts of UMI frequency were generated from FASTQ sequencing files by tabulating the number of occurrences of each 12nt sequence at the expected UMI position. The resulting count tables were further cleaned by requiring 1) the UMI matches the mixing code used to synthesize the library; 2) the UMI does not contain any ambiguous/uncalled bases; 3) the guide-UMI pair, or clone, is not a potential artifact of index bleed (the clone does not exactly match a more abundant clone on the same sequencing run); and 4) the clone is not a potential artifact of template switching (the UMI does not match a more abundant UMI paired with a different guide). To minimize the impact of singleton/low-frequency clones, a two-component mixture model fitted to the count frequency distribution and clones present in the low abundance component were dropped.

Clone counts present in tissues harvested at the end of the experiment (“endpoint” counts) were compared to clone counts present in the population of cells injected into the mice (“input” counts). To compute a robust gene-level enrichment score, guide-level clone counts were summed together for each gene prior to calling hits. All CRISPR screen analysis were performed using R (v4.2.3)[69].

For the genome-wide OT1/B16-Ova screen, gene-level log2 fold changes (endpoint over input) were computed using Limma-Voom as follows[70]. First, samples with low coverage (< 1,000 total clones and/or 0 clones recovered for > 50% of genes) were removed from the analysis. For the remaining samples, clone counts were summed at the gene level, and a pseudocount of 1 was added to each gene-level clone count. Genes that had fewer than 3 guides and/or low coverage (< 200 counts per million) in the input were removed from the analysis. Clone counts were normalized by dividing by the median clone count for each sample, and the resulting normalized counts from all sgRNA libraries were merged into one count matrix with four endpoint replicates. Genes screened in more than one library were summarized by calculating the mean normalized clone count across libraries. The EdgeR package (v3.40.2) in R was used to convert the counts table to a DGEList object and to calculate normalization factors using the trimmed mean of M-values (TMM) method[71]. The voom function from the limma package (v3.54.2) in R was then used to normalize the counts, model the mean-variance relationship, and calculate precision weights for each observation[72]. Using the standard limma empirical Bayes pipeline, the precision weights were incorporated into a linear model comparing endpoint vs. input to estimate log2 fold changes. Analysis based on read counts was performed with the same approach used for clone counts, except a pseudocount of 100 was added to each gene-level read count, and genes with < 100 counts per million were excluded. Gene set enrichment analysis was performed using the fgsea package (v1.24.0) in R[73]. After removing common essential genes, as determined by DepMap 23Q4 (downloaded from https://depmap.org/portal/data_page/), genes were ranked by signed p-values [-log10(p-value) * sign(logFC)] and analyzed for enrichment using the mouse Hallmark gene sets (v2022.1) from the Molecular Signatures Database[74–77].

To determine the ranking of individual guides for the genome-wide OT1/B16-Ova screen, guide-level log2 fold changes were calculated as follows. Samples that did not pass the quality control described in the gene-level analysis were removed, and a pseudocount of 0.5 was added to each guide-level clone count. Guides with low coverage (< 30 counts) in the input were removed from the analysis, and the remaining guide counts were normalized by dividing by the upper quartile clone count for each sample. Normalized counts for the different libraries were merged, and guides present in more than one library were summarized by calculating the mean normalized count across libraries. For each replicate, the log2 fold change of endpoint counts over input counts was calculated for each guide. The mean log2 fold change across replicates was calculated for each guide, and the resulting scores were centered to the median score.

Scores for the PMEL/MC38-gp100 screen were calculated as follows. First, samples with low coverage (< 100 total clones) were removed from the analysis. Endpoint replicates were then pooled together to boost the clone count for each gene, and a pseudocount of 1 was added to each gene-level clone count. For each library, the log2 fold change of endpoint counts over input counts was calculated for each guide and then centered to the median log2 fold change. Genes that had fewer than 7 guides and/or fewer than 5 endpoint clones after pooling replicates were removed from the analysis. The scores for each library were then combined, taking the mean score for genes present in more than one library. The same approach was used to calculate guide-level log2 fold changes, with an additional filtering step of removing guides with low coverage (< 10 counts) in the input.

For the lib55 OT1 screens performed with B16-Ova and EG7-Ova, gene-level log2 fold changes were computed using Limma-Voom as follows. To help boost clone counts for the EG7-Ova screen, which had slightly lower coverage than the B16-Ova screen, endpoint replicates were pooled together into 3 groups. Samples with low coverage (0 clones recovered for > 50% of genes) were removed from the analysis. Clone counts were then summed at the gene level, and a pseudocount of 1 was added to each gene-level clone count. Genes that had fewer than 5 guides and/or low coverage (< 1000 counts per million) in the input were removed from the analysis. Scores were then calculated with Limma-Voom, as described for the genome-wide OT1/B16-Ova screen. Guide-level log2 fold changes were calculated by first adding a pseudocount of 1 to each guide-level clone count and computing the log2 fold change of endpoint counts over input counts after removing guides with low counts in the input. For each replicate, log2 fold changes were centered to the median, and the resulting scores were averaged across replicates for each guide.

The combination sgRNA library contained 2 different guides targeting each individual gene, resulting in a total of 8 unique guide combinations for each gene pair, taking position into account. For the combination screens, the clone counts for each unique guide combination were summed together at the gene pair level after pooling endpoint replicates to help boost the signal. Using the olfactory receptor clone counts as a neutral control, enrichment scores were calculated by taking the logarithm of the ratio of the odds of enrichment for the gene pair of interest and the odds of enrichment for the neutral control:

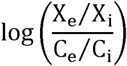

Where X_e_ and X_i_ are clone counts for the gene pair of interest in the endpoint and input samples, respectively, and C_e_ and C_i_ are clone counts for the olfactory receptor control paired with itself in the endpoint and input samples, respectively. Odds ratios and significance values were calculated in R using the fisher.test (alternative = ‘two.sided’) function. The log-odds ratio was calculated by taking the natural logarithm of the odds ratio + 0.1. The same approach was used to calculate guide-level scores after removing guides that were not sequenced in the input. To quantify combination effects, the difference in log-odds ratios between the observed and expected gene-level effects was calculated as follows:

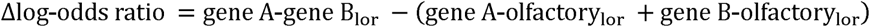

Where the observed effect was the measured log-odds ratio of gene A paired with gene B, and the expected effect was calculated as the sum of the log-odds ratios for gene A paired with the olfactory receptor control and gene B paired with the olfactory receptor control.

### Tissue Processing

For tissue processing of syngeneic tumor studies, single cell suspensions were generated from spleen, lymph nodes, blood and tumors. Spleens were processed using non-enzymatic digestion on GentleMacs protocol M spleen 01 01, RBC depleted with ACK (Lonza), filtered, washed, and resuspended in FACs buffer for staining. Lymph nodes were processed by gently pushing the tissue through a 35 µm filter, washing and resuspending in FACs buffer for staining. Blood samples were collected by either cardiac punction at termination or via tail vein at the indicated time points and were processed using ACK (Lonza) to deplete red blood cells. Tumor tissues were dissociated using mouse/human tumor dissociation kit (Miltenyi Biotec), wherein tumors are chopped into smaller pieces, incubated for 5-10 min at 37C in the enzyme mixture and then fully dissociated using tumor program 37C_TDK_1 as recommended by manufacturer (Miltenyi Biotec). Samples were resuspended at 10×10^6^ cells/mL in FACS buffer for staining or further purified using CD45 (TIL) MicroBeads (Miltenyi cat#130-110-618) isolation for single cell sequencing. Isolated CD45+ cells were then resuspended in PBS+0.4% BSA for sequencing.

### Flow Cytometry

Sample Processing: Single cell suspensions were processed on a 96 well v-bottom plate (Falcon, Cat#353077) for flow cytometry. Cells were washed with PBS (Gibco, 14190-136), stained with viability dye (Invitrogen, Cat#L34976), washed then and stained with a master mix of antibodies in Brilliant Stain Buffer (BD Biosciences, cat#566349) and Fc block (BD Pharmingen, Cat#553142) for surface staining. Following surface staining, cells were fixed by fixation buffer (BD Biosciences, 554655). For intracellular staining of transcription factors, samples were fixed and permeabilized using FoxP3 Fixation/Permeabilization kit (Invitrogen, 00-5523-00) and stained with master mix for transcription factors detection overnight. Samples were acquired on a BD FACS Symphony A3 or BD LSR-Fortessa and analyzed using Flowjo software (V10, Treestar).

### Editing Assessment by Amplicon Sequencing

Genomic DNA (gDNA) was prepared from cell pellets using the XTRACT 16+ gDNA cultured cells kit. Amplicons spanning the genomic on-target loci were amplified from gDNA by PCR using primer sets targeting the edit site. The resulting amplicons were sequenced using a NextSeq 500/550 Mid Output Reagent Cartridge v2 by 150 bp single-end NGS on a NextSeq500. Sequencing reads were computationally aligned to on-target sites, and the percentage of reads displaying an edited DNA sequence was determined. Target editing efficiencies for single-edit OT1s evaluated in Figure 3A were sgOlf: 56%; sgPD-1: 45%; sgSocs1: 88%; sgRegnase1: 82%; sgRoquin-1: 82% and sgIκBα: 71%. For dual-edit OT1s, editing efficiencies were sgSocs1 x Roquin-1: 85% and 74%; sgSocs1 x sgIκBα: 78% and 66%; and sgSocs1 x sgRegnase1: 80% and 75%. Target editing efficiencies for single-edit PMELs in Figure 3C were sgOlf: 69%; sgPD-1: 68%; sgSocs1: 93%; sgRegnase-1: 91%; sgRoquin-1: 81% and sgIκBα 87%. For dual-edit OT1s, editing efficiencies were sgSocs1 x Roquin-1: 90% and 79%; sgSocs1 x sgIκBα: 88% and 81%; and sgSocs1 x sgRegnase1: 82% and 89%. Target editing for OT1s in Figure 4 were sgOlf: 72%; sgPD-1: 65%; sgSocs1: 91%; sgRegnase-1: 85%; and sgSocs1 + sgRegnase-1: 91% and 85%, respectively.

### RNA-Seq

RNA was extracted using either Qiagen or Autogen kits as per the vendor’s guidelines. Libraries for RNA sequencing were prepared following the Truseq Stranded mRNA protocol. The final library concentrations were determined using the KAPA Library Quantification qPCR assay. Sequencing was carried out on a Nextseq2000 platform, aiming for a sequencing depth of 15 million reads for each sample.

Count and transcripts per million (TPM) data was generated by mapping the RNASeq fastqs to Gencode v43 for the human samples and Gencode m32 for the mouse samples using a custom version of the nf-core STAR-RSEM pipeline. The raw sequence image files from the sequencer were converted to the fastq format and checked for quality to ensure the sequencing quality scores did not deteriorate at the read-ends. Useful metrics such as the per base sequence quality, the per base sequence content, per base N content, sequence length distribution, adapter and k-mer content and sequence duplication levels were collected for each sequencing run using FastQC (Andrews S. FastQC: A quality control tool for high throughput sequence data. Http://Www.Bioinformatics.Babraham.Ac.Uk/Projects/Fastqc/. 2010). FastQCs were aligned to the human genome (GRCh38, Gencode version 43 or mouse genome Gencode m32) using STAR[78] aligner and viewed on Integrative Genome Viewer[79] (IGV). Only reads that were uniquely mapped were considered for read annotation to gene features. Reads were annotated with the human Gencode v43 or mouse Gencode m32 gene transfer format (gtf) file using RSEM.[80] RSEM provides both read counts and transcripts per million reads (TPMs) for each gene feature in the Gencode gtf. The R package DESeq2[81] was used for differential gene expression analysis. Genes were considered differentially expressed if they were UP-regulated by FC > +1.5 or DOWN-regulated by FC < - 1.5 with an adjusted p-value value < 0.05. For the PCA plots in Figure 6, the top 1000 variant genes (normalized, log2 transformed counts) were used after batch adjusting using ComBat from the sva package in R for RNA extraction kit, post-electroporation media and donor. Adjusting for donor was performed by regressing out donor with a linear model and using the residuals. Finally, the PC scores for PC1, PC2 and PC3 are plotted in a boxplot colored by group. Gene set enrichment analysis was performed using *fGSEA* in R. Briefly, genes were ranked by signed p-values [-log10(p-value) * sign(logFC)] for each edited TIL comparison with respect to its control and analyzed for enrichment using Hallmarks human or mouse gene sets from msigdb.

Transcription factor enrichment analysis was performed using *decoupleR*[82]. For each gene and comparison between edit vs control TILs, a linear univariate model was fitted. This model predicts the observed gene expression based solely on the TF-Gene interaction weights from the CollectTRI human TF network database. The sign of the t-value and the adj-p value of the model determines if the transcription factor is active.

### scRNA-Seq

Droplet-based 5’ single-cell RNA sequencing (scRNA-Seq) was performed using the 10x Genomics platform and libraries were prepared by the Chromium Single Cell 5’ Reagent kit according to the manufacturer’s protocol (10x Genomics, CA, USA). Cell Ranger (version 8.0.1) was used for gene expression quantification, TCR sequence assembly, and cell identification. RNA was aligned to the GRCm39-2024-A reference, and TCR sequences were aligned to the vdj_GRCm38_alts_ensembl-7.0.0 reference, both provided by 10x Genomics. Seurat (version 5.1.0)[83] was used to process feature-barcode matrices, align gene expression and TCR datasets, filter cells, identify and annotate clusters, and perform differential gene expression analyses.

Briefly, cells in which fewer than 200 unique genes were detected and cells where the fraction of counts belonging to mitochondrial genes (%mito) exceeded 10% were removed. In addition, cells were scored for how well they expressed S- and G2M-phase gene sets (Seurat) using CellCycleScoring, using murine equivalents of the human genes. For each cell, a value CC_difference_ was calculated as the difference between that cell’s S-phase and G2M-phase scores. Samples were processed using the scTransform workflow (Seurat), regressing %mito and CC_difference_. TCR variable and joining genes were removed from the list of genes used for principal component analysis and clustering to avoid clonotype-specific clusters. Dimensionality reduction was performed with 20 principal components, and downstream clustering was performed with resolutions typically between 0.5 and 1.0. Clusters were annotated based on expression of cell type-specific marker genes.

To create a T cell subset from the original CD45+ sample, only cells occupying T cell clusters based on expression of Cd3d with a corresponding TCR sequence were isolated and re-clustered. The final Cd8+ T cell subset was created from the T cell subset by combining 1) all OT-1 cells and 2) any endogenous T cells occupying Cd8+ clusters with equal-or-greater Cd8a expression than Cd4 expression. OT-1 cells were identified as cells containing only the CDR3 Tcrb sequence CASSRANYEQYF or CASSRANYEQYF paired with the CDR3 Tcra sequence CAASDNYQLIW. Differentially expressed genes were identified using the Wilcoxon Rank Sum test. Gene set scoring was performed on single cells using the rank-based method Singscore[84,85], and scores were averaged per cluster.

### TIL Culture

Viably cryopreserved tumor fragments were seeded into G-Rex 6 Well plate (5 fragments per well) in TIL complete media (10% HABS, 1X Pen/Strep, 10 mM HEPES, 1 mM Sodium Pyruvate, 5.5*10-5M 2-mercaptoethanol in RPMI1640) containing 6,000 IU/ml recombinant human IL-2, 10 µg/ml 4-1BB agonist antibody Urelumab (Wuxi Biologics), 30 ng/ml CD3 antibody OKT3 and cultured at 37°C with 5% CO2 for 11 days. During this process, termed pre-electroporation expansion (pre-EP), 6,000 IU/ml of recombinant human IL-2 was added every 3-4 days. Pre-EP TIL underwent CRISPR/Cas9 engineering (for KSQ-001EX, sgRegnase-1 or KSQ-004EX) or were left untreated (matching No EP controls). Briefly, pre-EP TIL were counted, centrifuged at 300 × g for 7 minutes and resuspended with MaxCyte electroporation buffer according to manufacturer protocol. Ribonucleoprotein (RNP) master mixture was added to the cell suspension at a final concentration of 2.4 µM u728 sgRNA, 2.4 µM j11d sgRNA and 2.08 µM Cas9 protein and transferred to processing assembly (MaxCyte). Cells were electroporated on a MaxCyte ExPERT electroporator using the “Optimization #9” program. After electroporation, cells were transferred to a recovery vessel and incubated at 37°C for 20 minutes for recovery. After recovery, cells were transferred to a G-Rex vessel (seeding density per vessel used was according to manufacturer recommendation) containing 50% TIL complete media + 50% AIMV for 10-12 days. During this process, termed post-electroporation expansion (post-EP), 3000 IU/mL of recombinant human IL-2 was added every 2-3 days. On Day 10 post-EP, cells were harvested and cryopreserved using 100% CryoStor10 at 50 × 10^6^cell/ml in a volume of 0.5 to 1 ml/vial. Cell pellets were also collected to determine editing efficiency by Next Generation Sequencing (NGS).

### In vitro Assays

Cytotoxicity and IFNγ Release Assay: A375-OKT3 cells engineered to express red fluorescent protein (RFP) were cultured in DMEM supplemented with 10% heat-inactivated FBS and 1× pen/strep solution. Three days before assay initiation, cells were harvested via TrypLE, with 10,000 cells/well plated in 100 μl RPMI supplemented with 10% heat-inactivated fetal bovine serum (FBS) and 1× pen/strep solution in ultra-low attachment U-bottom plates to allow for spheroid formation. The day prior to assay initiation, TIL were thawed and rested in 50% of TIL complete media + 50% AIMV supplemented with 100 U/mL IL-2. On the day of assay initiation, TIL were added at 1 10:1 effector to target (E:T) ratio to the spheroid plate. Images were taken via Incucyte S3 at 4× magnification in the red fluorescence, brightfield, and phase channels to monitor spheroid growth or regression. TIL cytotoxic activity was assessed at 72 hours by changes in red fluorescent intensity as a function of tumor spheroid cell growth or death, normalized to the first co-culture time point. Supernatant was collected 24 hours after the addition of TIL to target cells, and cytokine concentration was determined by MSD immunoassay following manufacturer protocol.

Chronic Stimulation Assay: A375 cells, expressing the high affinity membrane-bound OKT3 and red fluorescent protein (RFP), were cultured in DMEM supplemented with 10% heat-inactivated FBS and 1× pen/strep solution. The day before assay initiation, cells were harvested via TrypLE, with 3,000 cells/well plated in 100 μl TIL complete media in flat-bottom plates. On the day of assay initiation, TIL were thawed and added at a 2:1 effector to target (E:T) ratio to the A375-mOKT3, assuming doubling of the tumor cells in one day. Images were taken via Incucyte S3 at 20× magnification in the red fluorescence and phase channels to monitor tumor cell growth. Every 3-4 days, 50% of the cells were re-seeded in a new plate containing fresh A375-mOKT3 plated the day before. Throughout the experiment, the assay media was supplemented with 10 IU/mL of IL-2. At the end of each round, TIL were counted using CountBright Plus Absolute Counting Beads (Invitrogen), and proliferation of TIL was calculated.

## QUANTIFICATION AND STATISTICAL ANALYSIS

Statistical analyses were performed using GraphPad prism software or R using the indicated statistical test, with ns = no significance; * = p < 0.05; ** = p < 0.01; *** = p < 0.001 and **** = p < 0.0001.

