## Supplementary figures and images for "Dual-inactivation of Regnase-1 and SOCS1 rewires exhausted CD8^+^ T cell fate to enhance anti-tumor functionality"

### Graphic Abstract

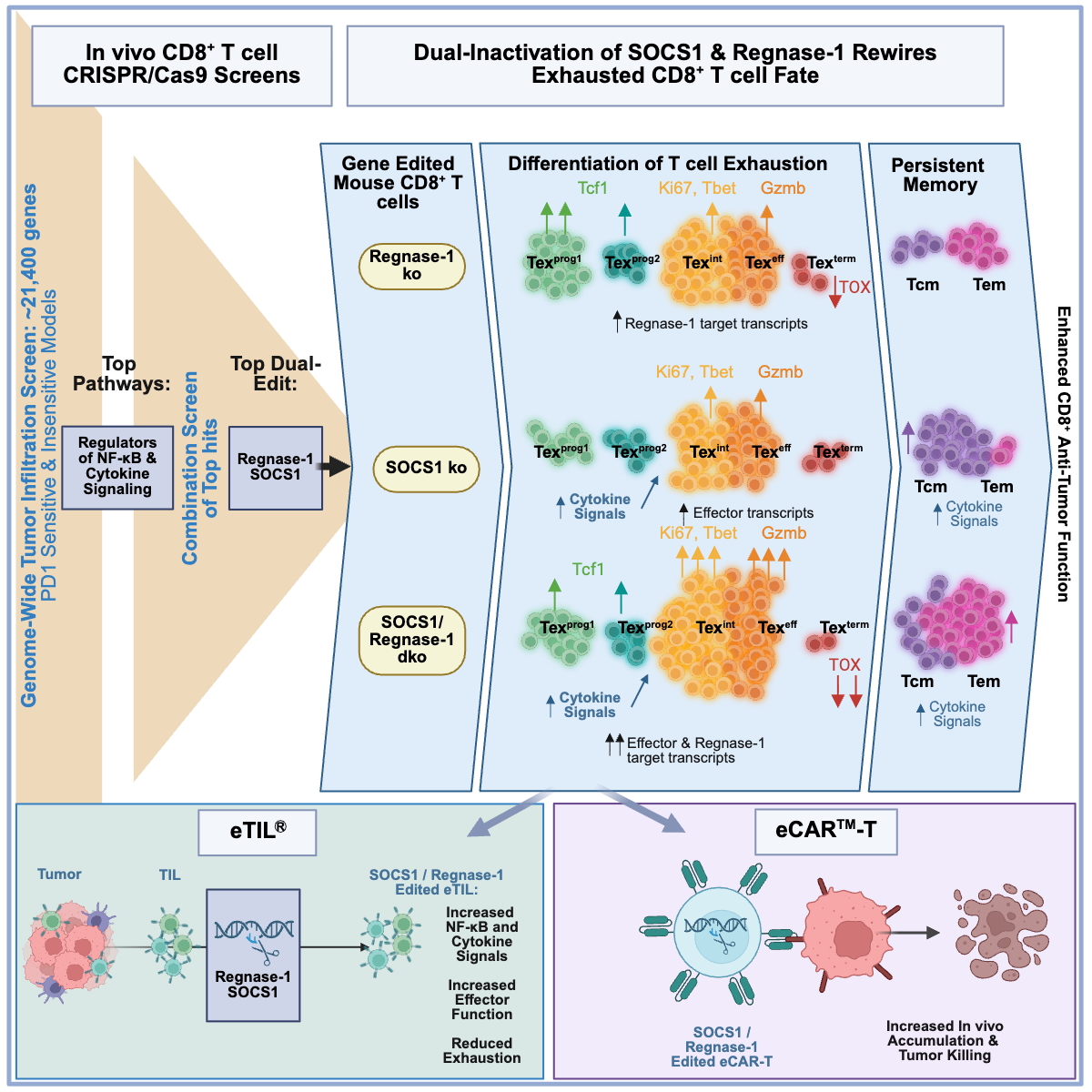
