## Supplemental Figures for "Dual-inactivation of Regnase-1 and SOCS1 rewires exhausted CD8^+^ T cell fate to enhance anti-tumor functionality"

### SUPPLEMENTARY INFORMATION CONTENTS


#### 1. SUPPLEMENTARY FIGURES

- **Figure S1:** Supporting data for Genome-wide CRISPR/Cas9 screen identifies top genes regulating the infiltration of transferred CD8 T cells into a B16-OVA solid tumor model
- **Figure S2:** The identification of top genes regulating the infiltration of transferred $CD8^+$ T cells into syngeneic tumor models
- **Figure S3:** Dual-sgRNA combination screens identify top target combinations regulating the infiltration of transferred $CD8^+$ T cells into murine syngeneic solid tumor models
- **Figure S4:** SOCS dual-edits drives durable OT1 persistence as $CD44^+CD62L^+$ $T_{cm}$ cells while Regnase-1 single and dual-edits drive OT1 persistence as $CD44^+CD62L^-$ $T_{em}$ cells
- **Figure S5:** Characterization and efficacy of SOCS1 and Regnase-1 single and dual-edited mTIL in a B16-OVA tumor model.
- **Figure S6:** Regnase-1 single and SOCS1/Regnase-1 dual inactivation in OT1s drives complete tumor regression and prolongs survival in a ‘large’ B16-OVA tumor setting and enhances the accumulation of $CD44^+CD62L^-$ cells in lymphoid tissues and tumor.
- **Figure S7:** scRNA-Seq on $CD45^+$ cells isolated from OT1s infiltrating TDLN and B16-OVA Tumors
- **Figure S8:** Regnase-1 single and SOCS1/Regnase-1 dual inactivation drives OT1 persistence as $CD44^+CD62L^-$ $T_{em}$ cells
- **Figure S9:** Differentially expressed proteins and genes in SOCS1 and Regnase-1 single and dual-edited Tex cells
- **Figure S10:** Characterization of KSQ-001EX, KSQ-004EX, sgRegnase-1 and No EP eTIL®
- **Figure S11:** Efficacy of SOCS1 + Regnase-1 CRISPR/Cas9-engineered meso eCAR™-Ts

#### 2. REFERENCES

### 1. Supplemental Figures

Supplementary Figure 1

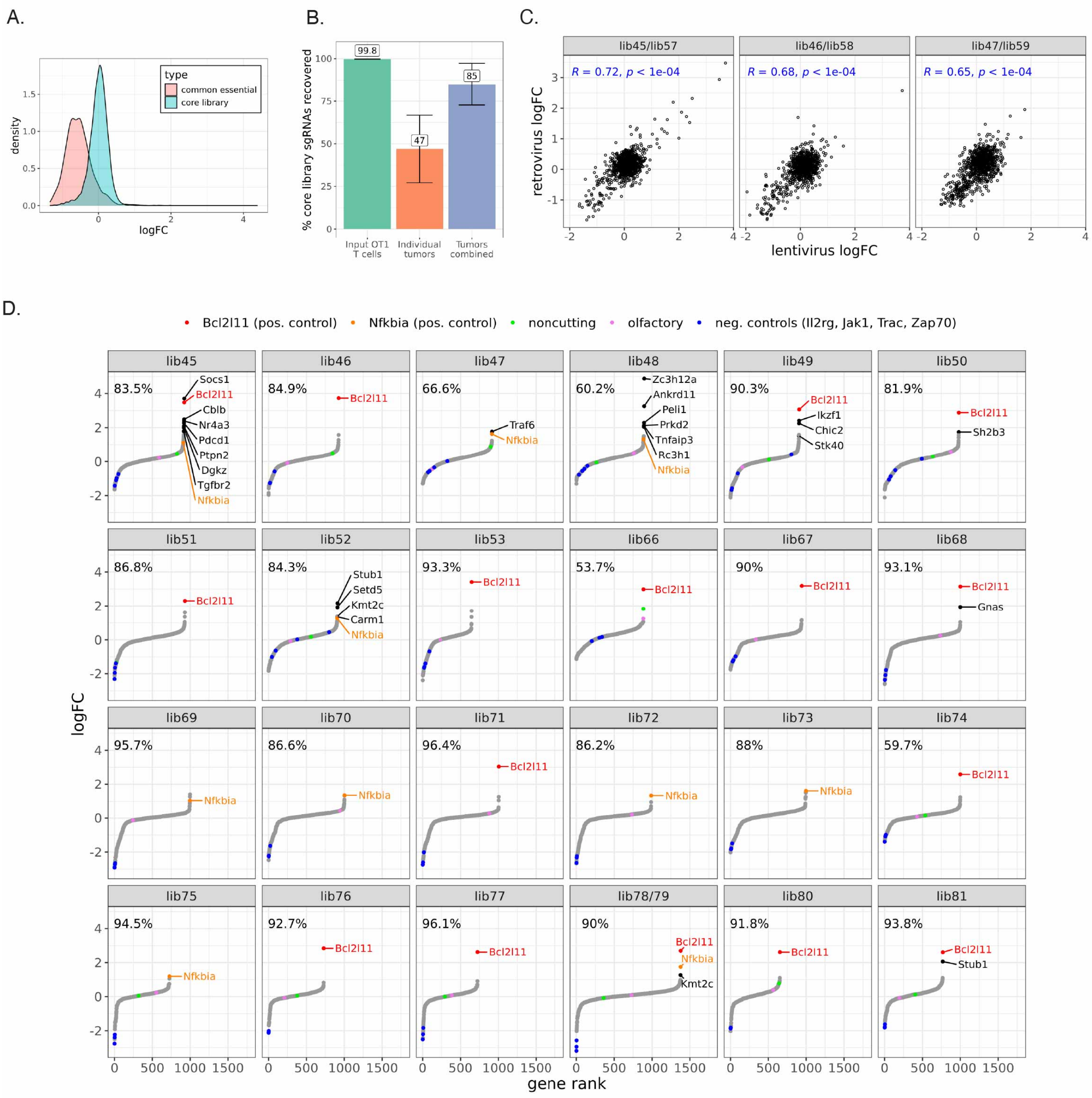

Supplementary Figure 1 (continued)

E.

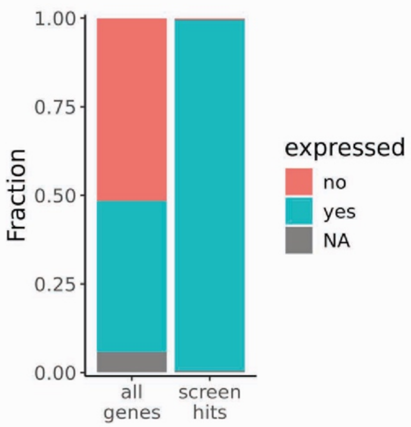


F.

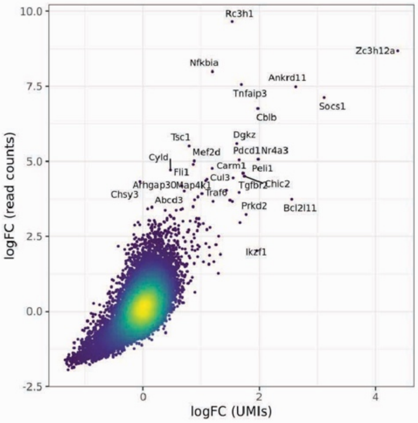


G.

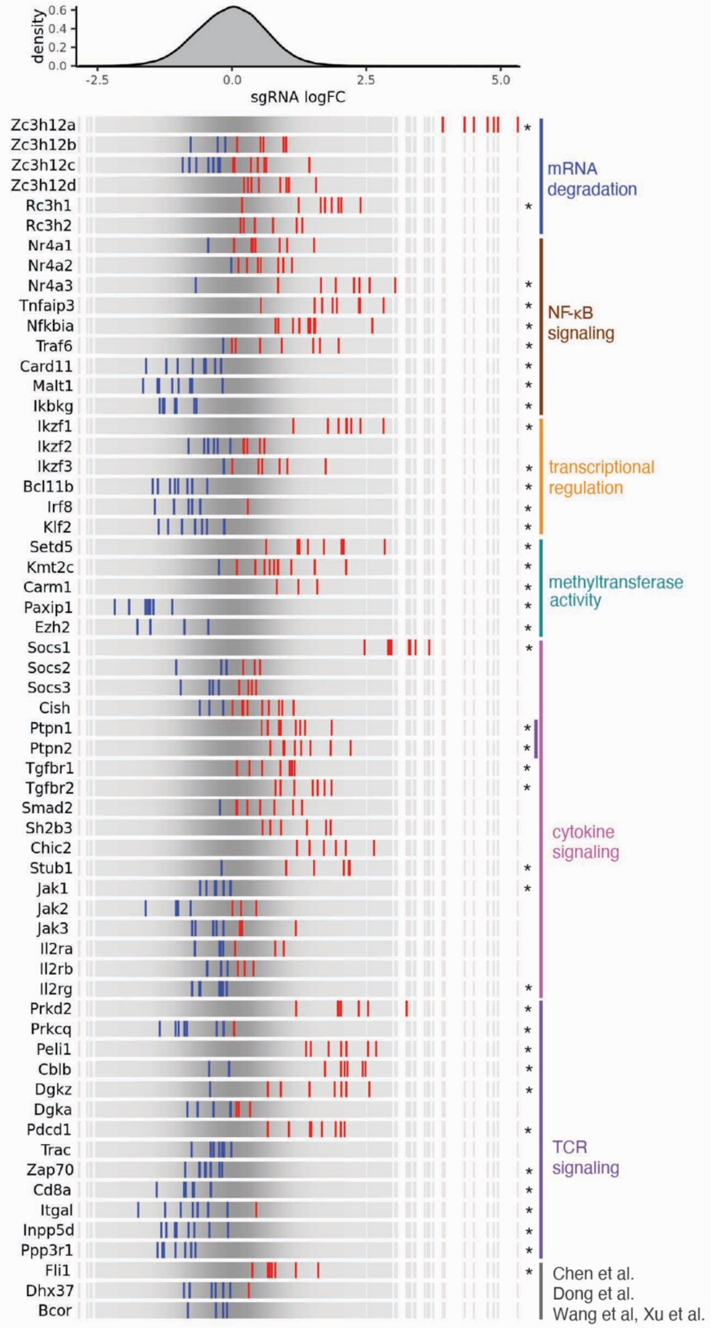


**Figure S1: Genome-wide CRISPR/Cas9 screen identifies top genes regulating the infiltration of transferred CD8 T cells into a B16-OVA solid tumor model**

Supporting data for Figure 1A-D. **(A)** Depletion of DepMap common essential genes (red). **(B)** sgRNA recovery from input samples (green; n = 24), individual tumor samples (orange; n = 96), and tumor samples pooled by bookshelf (blue; n = 24) across bookshelves. Bars represent the mean and error bars show the standard deviation. Only samples that were including in calling hits and that passed the quality filters described in the Supplementary Methods are shown. **(C)** Results from sgRNA library bookshelves screened using either lentivirus and positively selected using Thy1.1 (lib45, lib46, lib47) in comparison to the same sgRNA library bookshelf screened using retrovirus and positively selected using hCD2 (lib57, lib58, lib59). Pearson correlation coefficient (R) and p value (p) are shown in blue. **(D)** Rank plot by each sgRNA library bookshelf. Hits are depicted in black, controls shared across bookshelves as indicated, including positive controls (red, orange), noncutting (green), olfactory receptor (pink), and negative controls (blue). The frequency depicted in the upper left corner of each rank plot shows the percent of core library sgRNAs recovered in the endpoint after pooling replicates. **(E)** Expression status of 20,162 genes and screen hits only (n = 196) in the mouse T cell bulk RNA-Seq dataset described in Figure 3E. Screen hits = p value < 0.05 and abs(logFC) > 1, with expression cutoff of TPM > 1 across three independent RNA-Seq samples. NA = expression data not available. **(F)** Depiction of Figure 1B data as UMI vs Read-count analysis. **(G)** An expanded gene list from Figure 1C depicts enrichment (red) or depletion (blue) patterns of individual sgRNAs targeting top gene hits and controls by tumor infiltrating OT1s in comparison to input OT1s. Genes are clustered by established protein function, as indicated. Chen et al refers to a report describing Fli1 as a negative regulator of $CD8^+$ T cell accumulation in tumors[1] , Dong et al refers to a genome-wide screen describing Dhx37 as a negative regulator of $CD8^+$ T cell accumulation in tumors[2], Xu et al and Wang et all describe BCOR in combination with ZC3H12A as enhancing immortal $CD8^+$ T cells displaying stemness.[3,4] Asterisks denote genes with p values < 0.05.

### Supplementary Figure 2

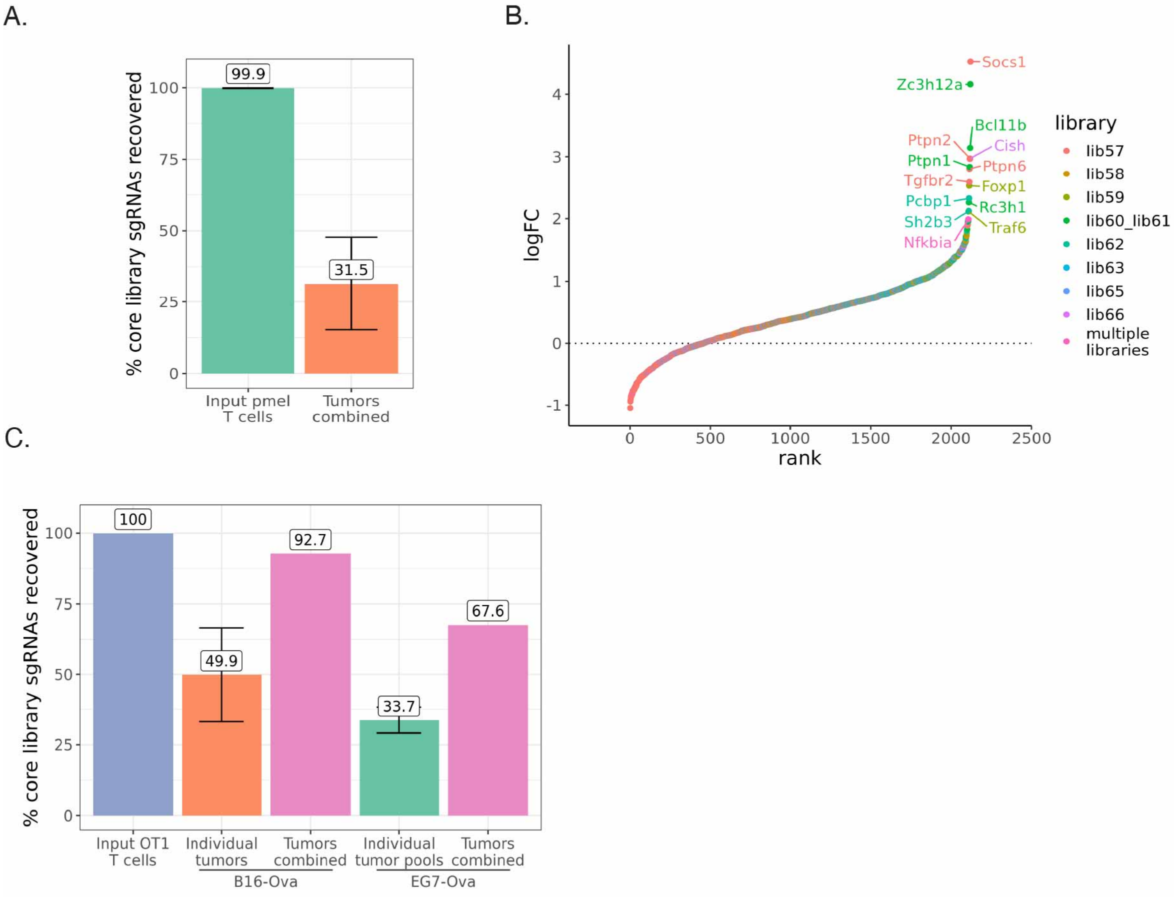


**Figure S2: Identification of top genes regulating the infiltration of transferred CD8⁺ T cells into syngeneic tumor models**

Supporting data for Figure 1E-F. **(A)** sgRNA recovery from input samples (green; n = 8) and tumor samples pooled by bookshelf (orange; n = 8) across bookshelves for the PMEL/ MC38-gp100 screen. Bars represent the mean and error bars show the standard deviation. Only samples that were including in calling hits and that passed the quality filters described in the Supplementary Methods are shown. **(B)** Rank plot colored by each sgRNA library bookshelf for the PMEL / MC38-gp100 screen. Only genes with at least 5 clones in the endpoint (after pooling replicates) are shown. **(C)** sgRNA recovery from the input sample (blue), individual B16-OVA tumor samples (orange; n = 5), individual EG7-OVA tumor pools (green; n = 3 pools of 2-3 replicates each), and all B16-OVA or EG7-OVA tumor samples combined (pink) for the OT1 Lib55 B16-OVA and EG7-OVA screens. Bars represent the mean and error bars show the standard deviation. Only samples that were including in calling hits and that passed the quality filters described in the Supplementary Methods are shown.

Supplementary Figure 3

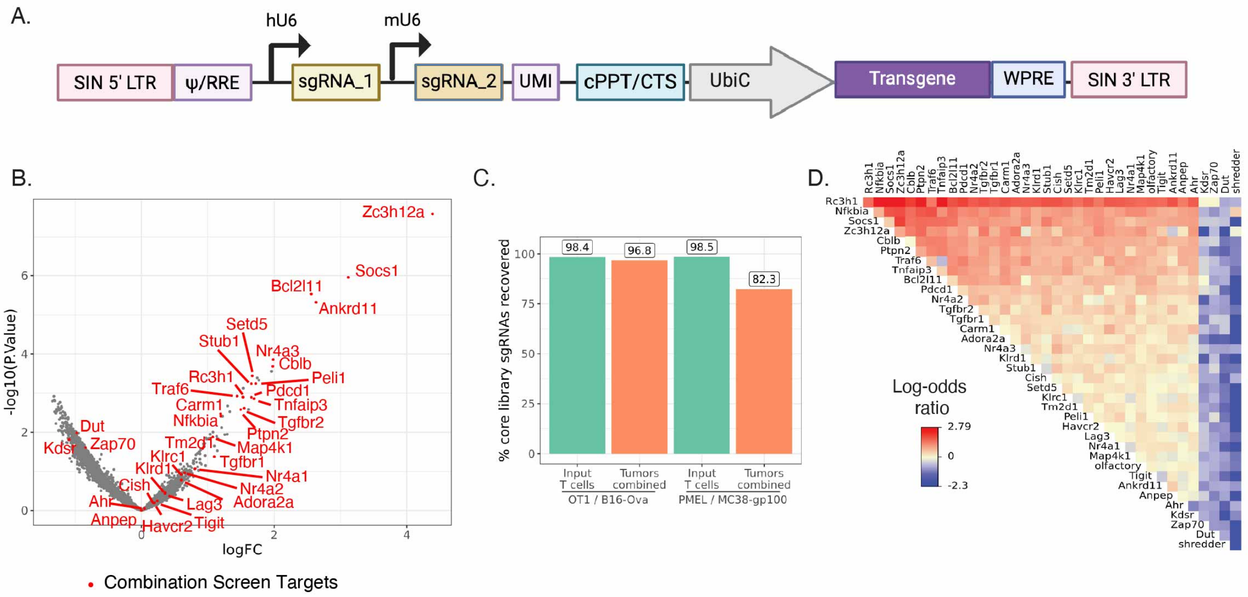


**Figure S3: Dual-sgRNA combination screens identify top target combinations regulating the infiltration of transferred CD8 T cells into murine syngeneic solid tumor models**

Supporting data for Figure 2. **(A)** Schematic of the combination screen plasmid design**. (B)** Selection of targets for the combination sgRNA library (highlighted in red) from the genome-wide OT1 / B16-OVA CRISPR/Cas9 screen in Figure 1B. **(C)** sgRNA recovery from the input samples (green) and pooled tumor samples (orange) for each depicted screen. **(D)** Results of a combination screen in the PMEL / MC38-gp100 model.

Supplementary Figure 4

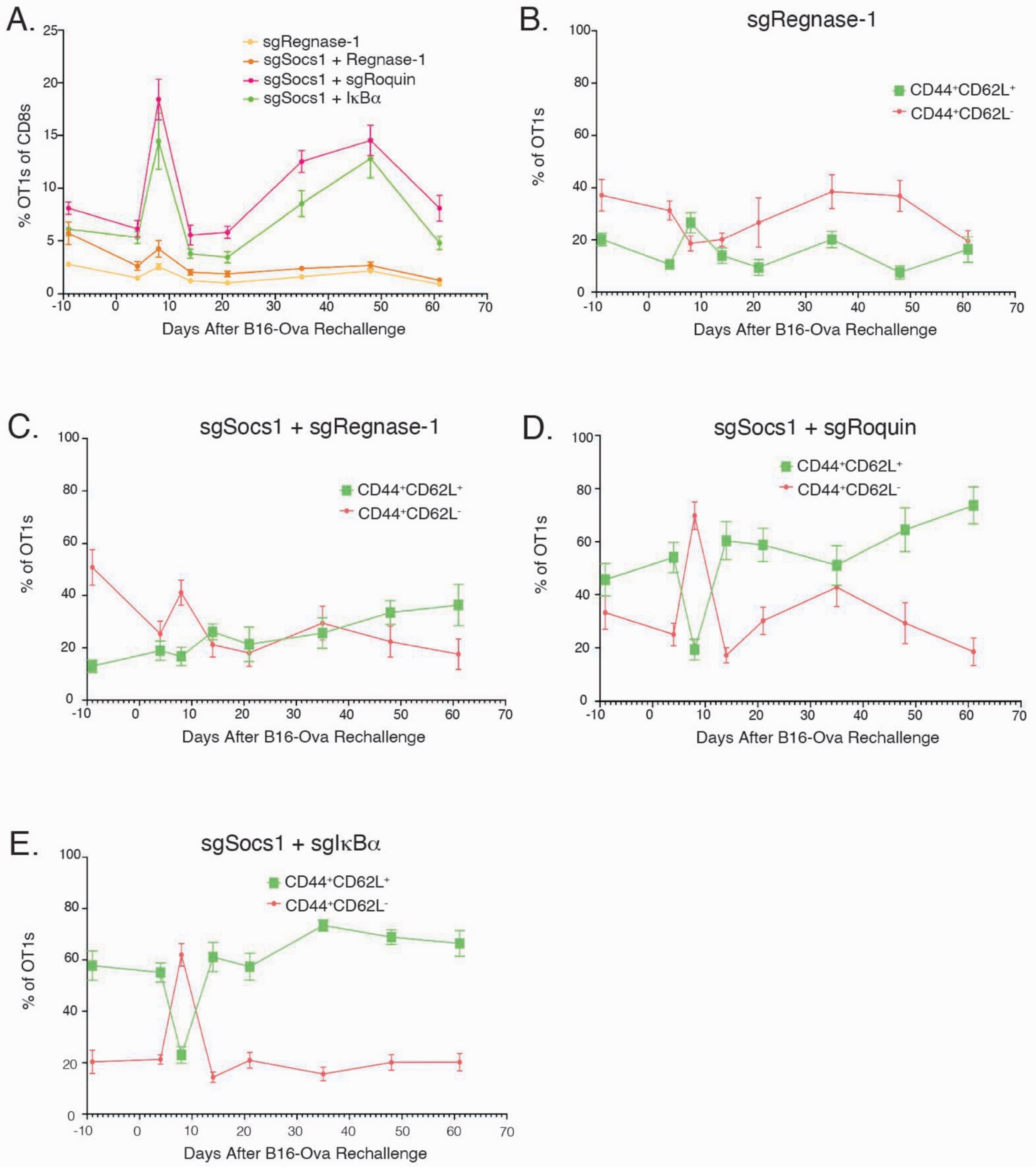


**Figure S4: SOCS dual-edits drives durable OT1 persistence as $CD44^{+}CD62L^{+}$ $T_{cm}$ cells while Regnase-1 single and dual-edits drive OT1 persistence as $CD44^{+}CD62L^{-}$ $T_{em}$ cells**

Supportive data for experiment depicted in Figure 3A-B. Regnase-1, SOCS1 + Regnase-1, SOCS1 + Roquin-1 and SOCS1 + Ikba single and dual-edited OT1-treated complete responder mice from Figure 3A were rechallenged with B16-OVA 78 days following initial T cell transfer as depicted in Figure 3B. **(A)** The frequency of single and dual-edited OT1s as peripheral blood $CD8^{+}$ T cells prior to and through day 61 following secondary B16-OVA re-challenge is depicted. Within single and dual-edited OT1s, the ratio of $CD44^{+}CD62L^{+}$ and $CD44^{+}CD62L^{-}$ cells within **(B)** Regnase-1 single edits, **(C)** SOCS1 + Regnase-1 dual-edits, **(D)** SOCS1 + Roquin-1 dual edits and **(E)** SOCS1 + IκBα dual edits are depicted.

Supplementary Figure 5

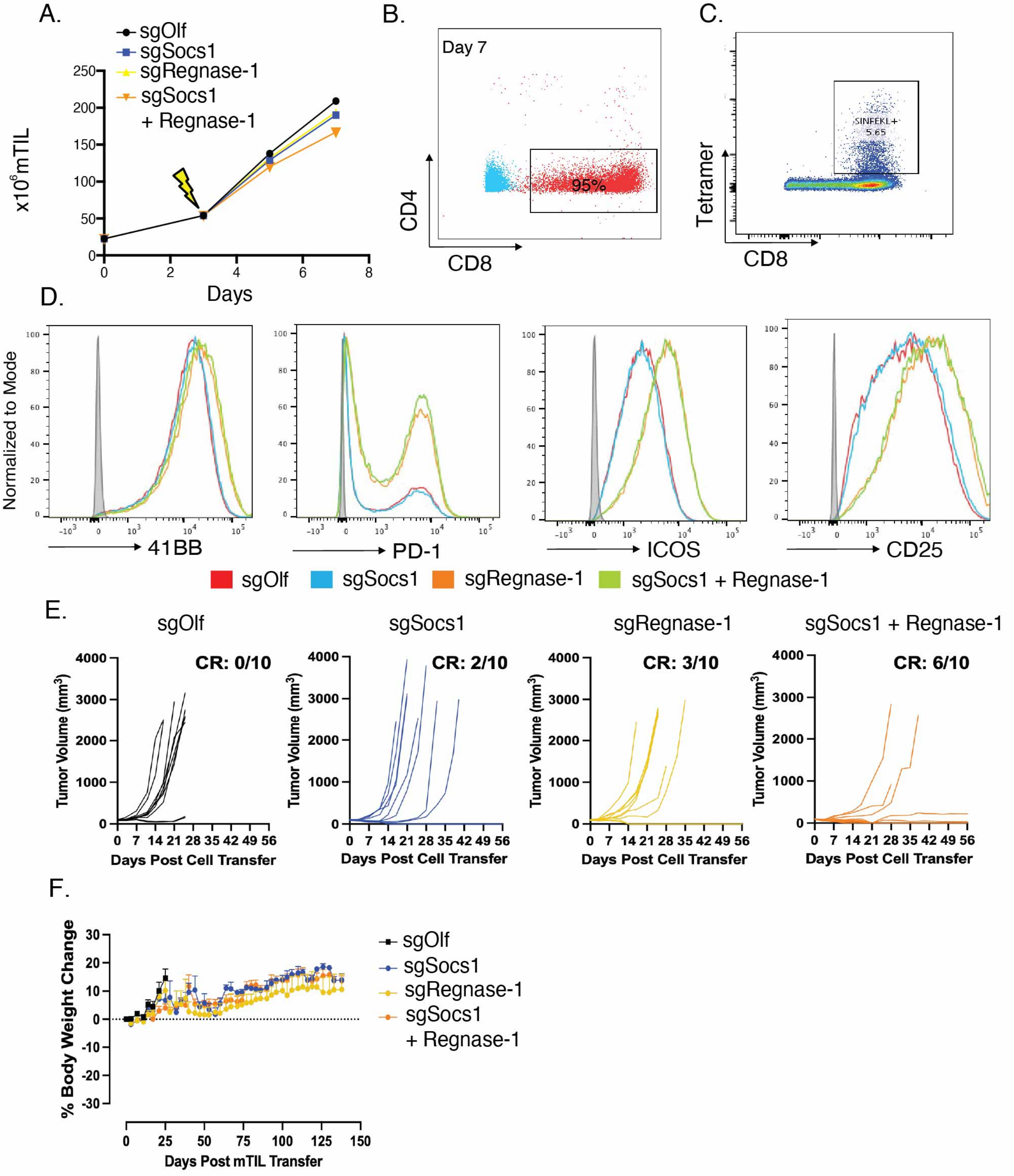


**Figure S5: Characterization and efficacy of SOCS1 and Regnase-1 single and dual-edited mTIL in a B16-OVA tumor model**

Supportive data for experiment depicted in Figure 3E-H. **(A)** Expansion of single and dual-edited mTIL over 7 days ex vivo culture and electroporation. **(B)** Following expansion, mTIL cellularity was characterized. **(C)** Specificity of mTIL for B16-OVA was determined by SIINFEKL tetramer staining. **(D)** Cell surface expression of 4-1BB, PD-1, ICOS and CD25 by single and dual-edited mTIL was determined by flow cytometry, with gray histogram indicating FMO controls. **(E)** Spider plots of individual mouse B16-OVA tumor growth curves and number of complete responders depicted in mTIL-treated mice. Same experiment as in Figure 3F. **(F)** Body weight over time by mTIL treatment group, as indicated. For survival analysis, Log-rank (Mantel-Cox) test was used (*** = p value <0.001).

Supplementary Figure 6

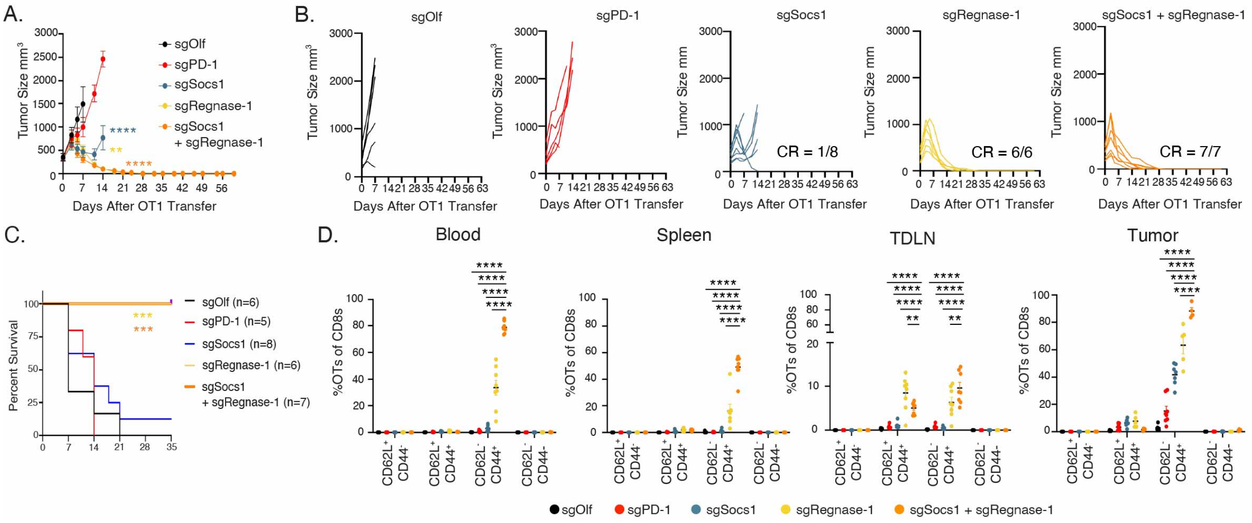


**Figure S6: Regnase-1 single and SOCS1/Regnase-1 dual inactivation in OT1s drives complete tumor regression and prolongs survival in a 'large' B16-OVA tumor setting and enhances the accumulation of CD44⁺CD62L⁻ cells in lymphoid tissues and tumor**

Supportive data for experiment depicted in Figure 4. **(A)** Tumor growth curves of mice bearing 'large' 340mm$^3$ B16-OVA tumors with single or dual-edited OT1 cells, as indicated. **(B)** Spider plots of individual tumor growth curves and number of compete responders (CRs) by single and dual-edited OT1s. sgSocs1: 1/8 CRs; sgRegnase-1: 6/6 CRs, and sgSocs1 + sgRegnase-1 with 7/7 CRs, as indicated. **(C)** Overall survival benefit by treatment group. **(D)** Frequencies of CD62L$^+$CD44$^+$, CD62L$^+$CD44$^-$, CD62L$^-$CD44$^-$ and CD62L$^-$CD44$^+$ OT1s of total CD8$^+$ T cells was determined in blood, spleen, TDLNs, and tumor between treatment groups. To evaluate statistical significance in tumor growth in Figure S6A, a 2-way ANOVA with Tukey's multiple comparisons was utilized. For survival analysis in Figure S6C, Log-rank (Mantel-Cox) test was used. Results of a 2-way ANOVA was used determine statistical significance between treatment groups in Figure S6D, with ns = no significance, * = p value < 0.05, ** = p value < 0.01, *** = p value <0.001 and **** = p value < 0.0001.

Supplementary Figure 7

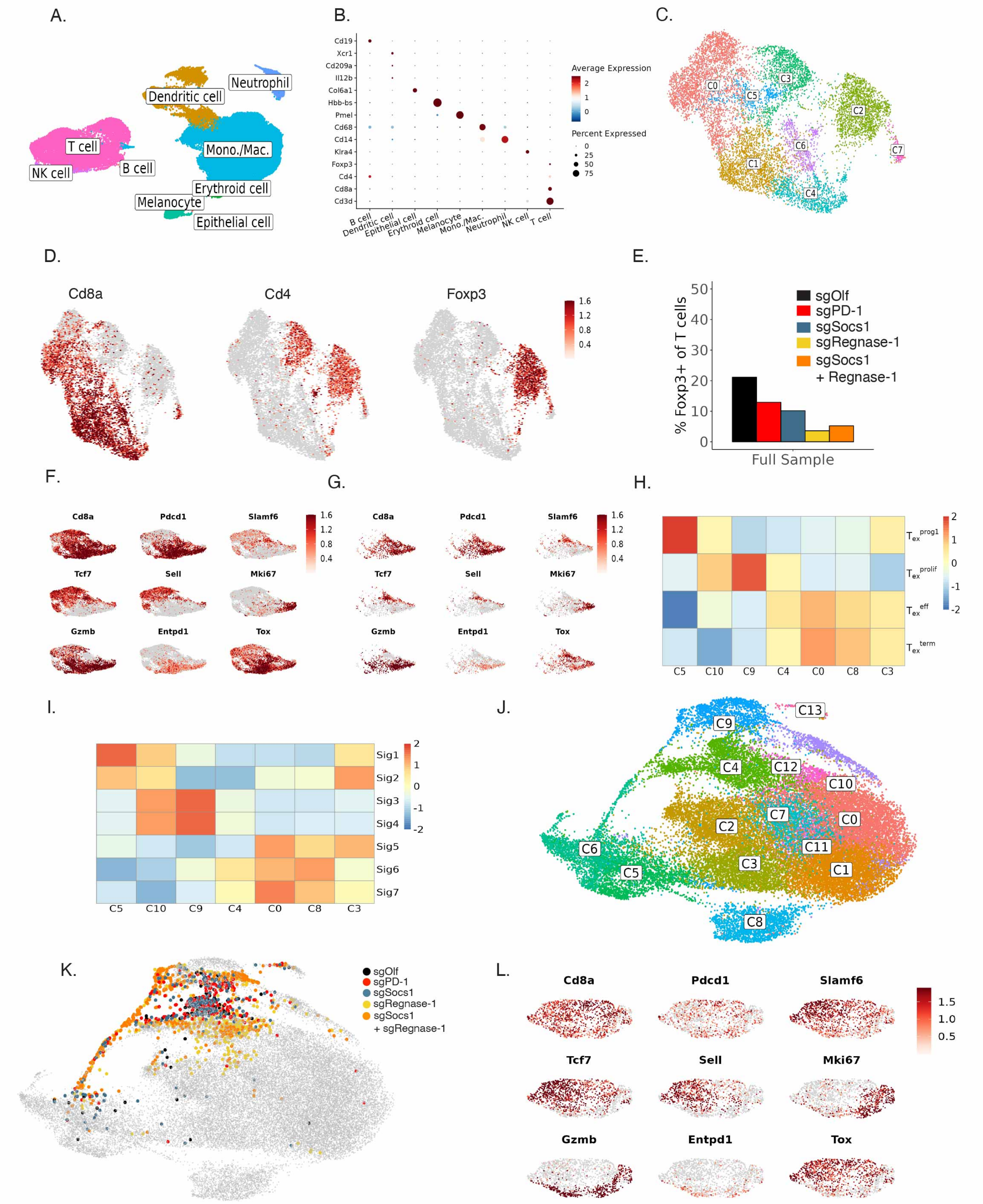


**Figure S7: scRNA-Seq on CD45⁺ cells isolated from OT1s infiltrating TDLN and B16-OVA Tumors**
Supportive data for experiment depicted in Figure 4. **(A)** General distribution of lymphocyte populations from CD45⁺ cells isolated from tumor annotated based on cell lineage-defining transcripts displayed by Dot

plot in **(B)**. **(C)** UMAP visualization of CD3$^+$ T cell clusters. **(D)** Expression of *Cd8a* (left), *Cd4* (center) and *Foxp3* (right) transcripts projected on CD3$^+$ clusters from Figure S7C. **(E)** Frequency of Foxp3$^+$ cells among total CD3$^+$ T cells by treatment group is depicted. **(F)** Expression of the indicated lineage-defining Tex transcripts in a UMAP visualization among total CD8$^+$ cells. **(G)** Expression of Tex-defining transcripts in a UMAP visualization limited to just OT1s. **(H)** Heatmap of median Singscores from Miller et al[5] and **(I)** Beltra et al[6] Tex gene signatures per cluster from Figure 4D. **(J)** UMAP visualization of CD8$^+$ clusters from TDLNs. **(K)** OT1 T cells overlayed on non-OT1 cells (gray cells) in TDLN, colored by treatment group, per legend. **(L)** Expression of indicated lineage-defining Tex transcripts in a UMAP visualization of re-clustered OT1s in TDLN.

Supplementary Figure 8

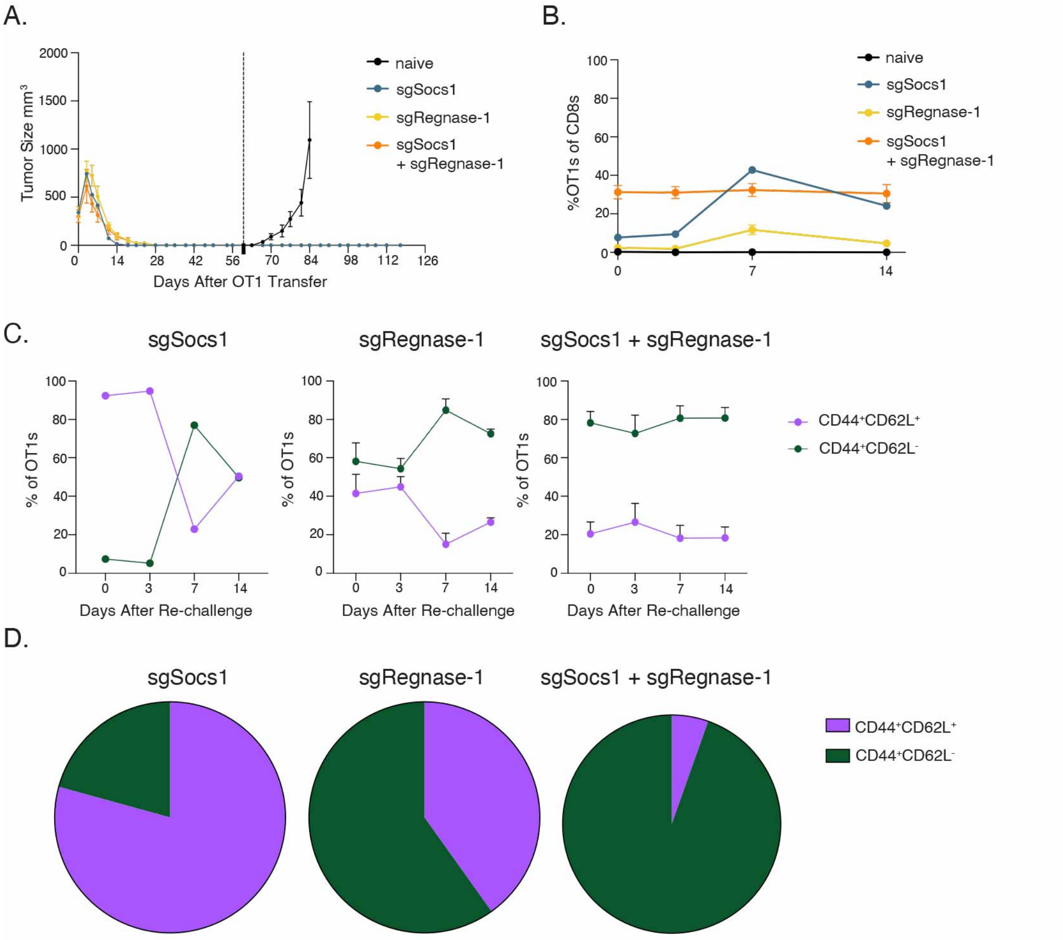


**Figure S8: Regnase-1 single and SOCS1/Regnase-1 dual inactivation drives OT1 persistence as CD44$^+$CD62L$^-$ T$_{em}$ cells**

**(A)** Mice undergoing complete tumor rejection in Figure S6A were re-challenged with B16-OVA tumor cells 60 days following initial T cell transfer, with naïve mice included as controls. Tumor growth by treatment group is depicted. **(B)** The frequency of OT1s as peripheral blood CD8$^+$ T cells prior to and following secondary B16-OVA re-challenge is depicted by treatment group. **(C)** Relative frequencies over time post-challenge of CD44$^+$CD62L$^+$ and CD44$^+$CD62L$^-$ cells among single and dual-edited OT1s. **(D)** The frequency of CD44$^+$CD62L$^+$ and CD44$^+$CD62L$^-$ cells within each treatment group 117 days following initial OT1 transfer.

Figure S9

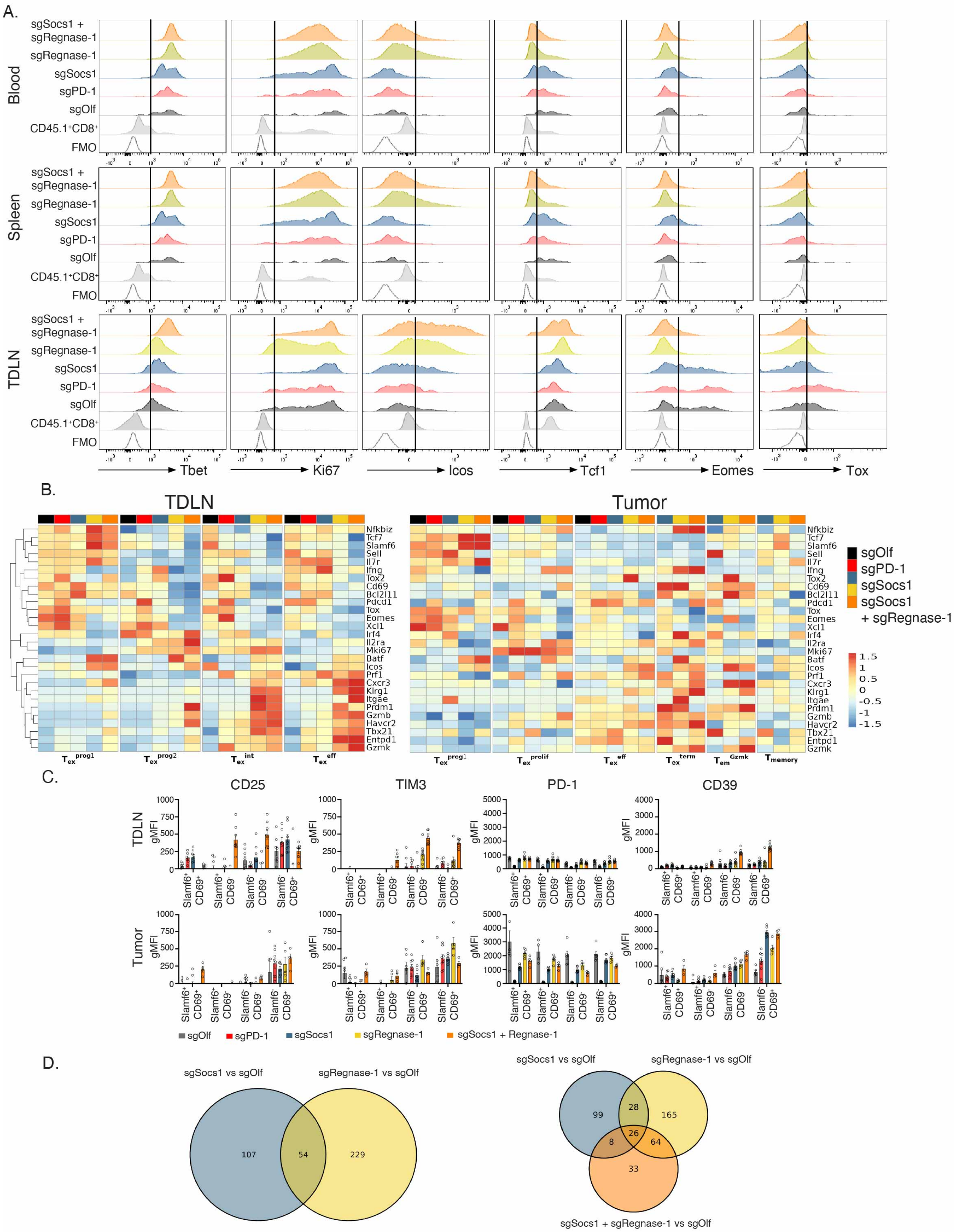
A.
Blood
Spleen
TDLN
sgSocs1 + sgRegnase-1
sgRegnase-1
sgSocs1
sgPD-1
sgOlf
CD45.1+CD8+
FMO
Tbet
Ki67
Icos
Tcf1
Eomes
Tox
B.
TDLN
Tumor
sgOlf
sgPD-1
sgSocs1
sgSocs1 + sgRegnase-1
Nfkbiz
Tcf7
Slamf6
Sell
Il7r
Ifng
Tox2
Cd69
Bcl2l11
Pdcd1
Tox
Eomes
Xcl1
Irf4
Il2ra
Mki67
Batf
Icos
Prf1
Cxcr3
Klrg1
Itgae
Prdm1
Gzmb
Havcr2
Tbx21
Entpd1
Gzmk
Tex prog1
Tex prog2
Tex int
Tex eff
Tex prolif
Tex term
Tem Gzmk
Tmemory
C.
CD25
TIM3
PD-1
CD39
gMFI
Slamf6+ CD69+
Slamf6+ CD69-
Slamf6- CD69-
Slamf6- CD69+
sgOlf
sgPD-1
sgSocs1
sgRegnase-1
sgSocs1 + Regnase-1
D.
sgSocs1 vs sgOlf
sgRegnase-1 vs sgOlf
107
54
229
99
28
165
8
26
64
33
sgSocs1 + sgRegnase-1 vs sgOlf

**Figure S9: Differentially expressed proteins and genes in SOCS1 and Regnase-1 single and dual-edited Tex cells**

Supportive data for experiment depicted in Figure 5. **(A)** Histograms depicting expression of the indicated proteins by flow cytometry in blood, TDLN and spleen OT1s by treatment group as indicated. **(B)** scRNA-Seq analysis of TDLN and tumor OT1s by Tex and memory cell subset. Expression of Tex lineage transcripts are depicted by treatment group. **(C)** Impact of single and dual-inactivation on the expression of indicated cell-surface receptor by FACS in TDLN and Tumor-derived OT1s by Slamf6$^{+}$CD69$^{+}$, Slamf6$^{+}$CD69$^{-}$, Slamf6$^{-}$CD69$^{-}$ and Slamf6$^{-}$CD69$^{+}$ subsets as indicated. **(D)** Venn diagram depiction of common DEGs between indicated SOCS1 and Regnase-1 single and dual-edited OT1 group comparisons with controls.

Supplementary Figure 10

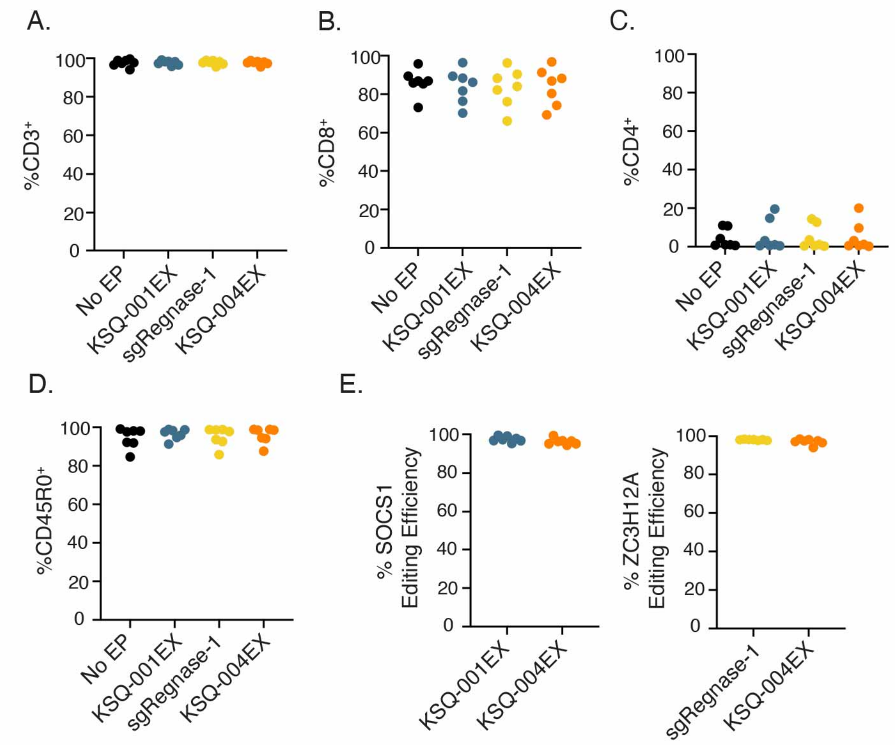


**Figure S10: Characterization of KSQ-001EX, KSQ-004EX, sgRegnase-1 and No EP TIL**
Supportive data for experiment depicted in Figure 6. No EP, KSQ-001EX, sgRegnase-1 and KSQ-004EX were evaluated by flow cytometry for expression of **(A)** CD3, **(B)** CD8, **(C)** CD4 and **(D)** CD45RO. **(E)** Editing efficiency of the *SOCS1* and *ZC3H12A* genomic cut-site by CRISPR/Cas9 RNPs as determined by AMP-Seq.

Supplementary Figure 11

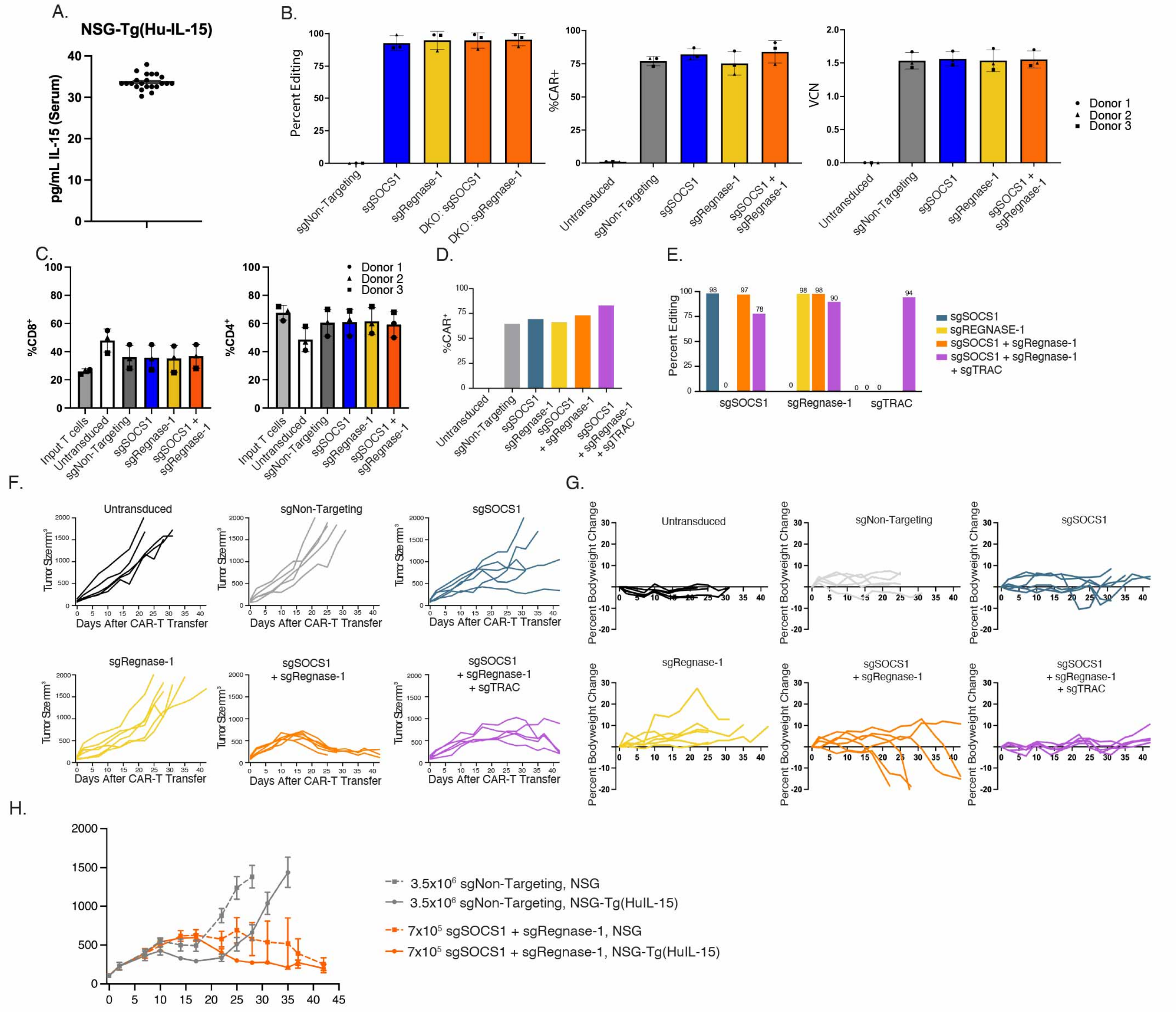


**Figure S11: Efficacy of SOCS1 / Regnase-1 CRISPR/Cas9-engineered meso eCAR-Ts**

Supportive data for experiment depicted in Figure 7. **(A)** Serum IL-15 levels from the NSG-Tg(Hu-IL-15) (Jackson Laboratories) mouse strain. Control, single and dual-edited meso eCAR-Ts were manufactured from three independent donors, with **(B)** editing efficiency of the *SOCS1* and *ZC3H12A* genes (left), meso CAR expression levels (middle) and vector copy number (VCN; right) quantified per donor and treatment group as depicted. CAR expression was determined by detecting RQR8 expression by flow cytometry. **(C)** meso CAR expression levels and **(D)** editing efficiency of the *SOCS1, ZC3H12A* and *TRAC* genes of donor used for in vivo efficacy. **(F)** Tumor growth spider plots depicted data from Figure 7D. **(G)** Body weight over time following meso eCAR-T transfer, per treatment group. **(H)** Efficacy of the of indicated dose of sgNon-Targeting or sgSOCS1 + sgRegnase-1 meso eCAR-Ts in the NSG or NSG-Tg(Hu-IL-15) mouse strains, as indicated.
